# Single-cell monitoring of dry mass and dry density reveals exocytosis of cellular dry contents in mitosis

**DOI:** 10.1101/2021.12.30.474524

**Authors:** Teemu P. Miettinen, Kevin S. Ly, Alice Lam, Scott R. Manalis

## Abstract

Cell mass and composition change with cell cycle progression. Our previous work characterized buoyant mass accumulation dynamics in mitosis (Miettinen et al., 2019), but how dry mass and cell composition change in mitosis has remained unclear. To better understand mitotic cell growth and compositional changes, we develop a single-cell approach for monitoring dry mass and the density of that dry mass every ∼75 seconds with 1.3% and 0.3% measurement precision, respectively. We find that suspension grown mammalian cells lose dry mass and increase dry density following mitotic entry. These changes display large, non-genetic cell-to-cell variability, and the changes are reversed at metaphase-anaphase transition, after which dry mass continues accumulating. The change in dry density causes buoyant and dry mass to differ specifically in early mitosis, thus reconciling existing literature on mitotic cell growth. Mechanistically, the dry composition changes do not require mitotic cell swelling or elongation. Instead, cells in early mitosis increase lysosomal exocytosis, and inhibition of exocytosis prevents the dry composition from changing. Overall, our work provides a new approach for monitoring single-cell dry mass and composition and reveals that mitosis is coupled to extensive exocytosis-mediated secretion of cellular contents.

## INTRODUCTION

Cells coordinate the activity of biosynthetic and catabolic pathways as well as the uptake and secretion of components in order to maintain appropriate molecular composition. In continuously proliferating cells, these processes are coordinated so that all cellular components are doubled during each cell cycle, whereas differentiating cells may change their composition to match their functions (Cadart et al., 2019; Lloyd, 2013; Miettinen et al., 2017; Schmoller and Skotheim, 2015). Changes in cells’ molecular composition are typically studied with approaches such as mass spectrometry, which can quantify the molecular details of a cell population after lysis. However, there is a lack of methods that can monitor the cellular composition of live single cells with high resolution, and thereby enable the study of dynamic and temporary events such as mitosis.

Cells’ total molecular content, i.e. dry mass, can be monitored using methods such as quantitative phase microscopy (QPM). Dry mass measurements are critical for understanding cell size and growth regulation, but they are limited in resolution and, more importantly, they do not inform us about the composition of the dry mass. One approach that overcomes this, is stimulated Raman scattering microscopy, which can measure the total amount of proteins and lipids in a live cell (Figueroa et al., 2021; Oh et al., 2020). While Raman scattering-based methods provide spatial resolution and details about the molecular changes taking place, they can also cause phototoxicity, which significantly limits the temporal resolution when tracking the same cell(s) over time (Zhang et al., 2021). Alternatively, QPM-based dry mass measurements can be coupled to microscopy-based total volume measurements to quantify dry density, an indicator of cell composition (Cooper et al., 2013; Odermatt et al., 2021; Venkova et al., 2021; Zlotek-Zlotkiewicz et al., 2015). However, these approaches include intracellular water content in the volume quantification, thereby making the approach mainly sensitive towards total molecular concentrations inside the cell.

Here, we introduce a new approach for monitoring single cell’s dry composition by repeatedly measuring the same cell as it grows. Cell’s dry mass and the density of that dry mass can be quantified by comparing the cell’s buoyant mass in normal water (H_2_O) and heavy water (deuterium oxide, D_2_O) -based solutions (Delgado et al., 2013). However, high concentration of D_2_O prevents cell proliferation and can result in cell death (Schroeter et al., 1992; Takeda et al., 1998). Here, we show that we can expose the cell to D_2_O only periodically (every 75 s) and for very short durations (10-15 seconds), thus allowing us to non-invasively monitor the dry mass and dry density of the same cell over many generations without compromising cell growth and viability. Importantly, this approach measures dry density independently of the cell’s water content and is therefore sensitive towards changes in the relative molecular (dry) composition, not the overall concentration of dry mass. For example, a decrease in the amount of cellular lipids increases the density of the cell’s dry mass, as lipids are lower in density than most other cellular components.

We have previously monitored the buoyant mass accumulation of single mammalian cells in mitosis (Miettinen et al., 2019) using the suspended microchannel resonator (SMR), a cantilever-based single-cell buoyant mass sensor (Burg et al., 2007). We found that buoyant mass accumulation rate is high from prophase to metaphase-anaphase transition, i.e. early mitosis. In contrast, studies using QPM have suggested that dry mass does not accumulate in early mitosis (Zlotek-Zlotkiewicz et al., 2015), or that cells may even experience a decrease in dry mass in early mitosis (Liu et al., 2020). If cells actually lose dry mass in mitosis, this would indicate previously unrecognized compositional changes taking place in mitosis. Although the exact dynamics of dry mass accumulation in mitosis remain debatable due to the limited resolution of existing approaches, our study addresses this seeming contradiction between buoyant and dry mass measurements by providing a high-resolution view of the mitotic dry mass and composition changes.

## MEASUREMENT METHOD

### Measurement principle

In our method, the cell first travels through the SMR while filled with H_2_O-based solution to obtain a buoyant mass measurement (Figure 1A). On the other side of the SMR, the cell is briefly exposed to D_2_O-based solution. Cells normally exchange their water content in ∼1 second (Potma et al., 2001), causing intracellular H_2_O to be exchanged to D_2_O. The cell then travels back through the SMR in D_2_O solution to obtain a second buoyant mass measurement (Figure 1B). As both of the buoyant mass measurements are carried out in solutions where the intracellular water content is neutrally buoyant with the extracellular solution, these two measurements can be used to calculate dry mass, dry density and dry volume of the cell (Figure 1C) (Delgado et al., 2013). Mathematically, this is can be written out as:

**Figure 1.**
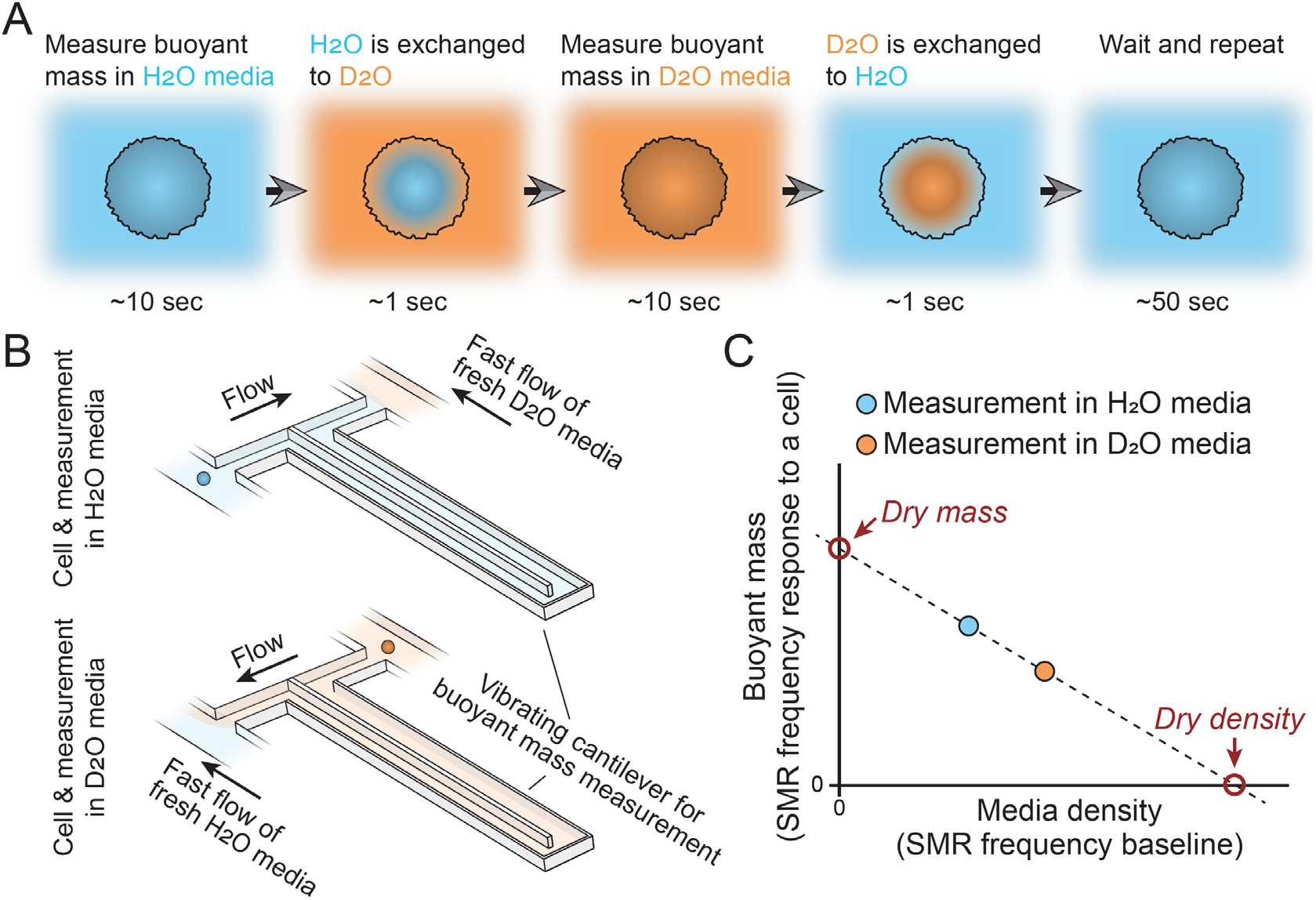
Schematic of dry mass and dry density measurements using the SMR. **(A)** The cell’s buoyant mass is first measured in normal, H_2_O-based culture media (blue), after which the cell mixes with D_2_O-based culture media (orange). The water content inside the cell exchanges to match the external water content. The cell’s buoyant mass is then measured again in the D_2_O-based media and the cell is mixed with normal, H_2_O-based media, where the cell waits for the next measurement. **(B)** In practice, these measurements are carried out using a suspended microchannel resonator (SMR), where H_2_O and D_2_O-based medias are kept on different sides of the cantilever. Continuous flushing of fresh media into the system prevents the two fluids from equilibrating over time. The cell is depicted as a small blue/orange sphere. **(C)** To calculate the dry mass and dry density of the cell, the two buoyant mass measurements are correlated as a function of media density. The SMR’s baseline signal (vibration frequency) is proportional to media density and this measurement is obtained immediately before and after each buoyant mass measurement (frequency response to a cell flowing through the SMR) to account for variability in media mixing.

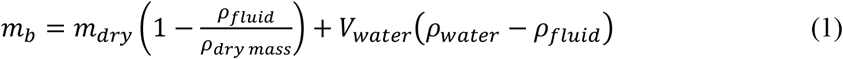

where *m*_*b*_ is the buoyant mass of the cell in a given fluid, *m*_*dry*_ is the dry mass of the cell, *ρ*_*fluid*_ is the density of the fluid outside the cell, *ρ*_*dry mas*_ is the density of cell’s dry mass, *V*_*water*_ is the volume of intracellular water content, and *ρ*_*water*_ is the density of intracellular water content. The first term of *equation 1* describes the contribution of the cell’s dry mass to buoyant mass and the latter term describes the contribution of the cell’s water content to buoyant mass. The densities of the extracellular fluid are not constant due to fluid mixing between measurements, but this is accounted for by directly measuring the extracellular solution density, i.e. the baseline frequency of the SMR, together with every buoyant mass measurement. As D_2_O and D_2_O-based media are very similar in density (1.10 g/ml), as are H_2_O and H_2_O-based media (1.00 g/ml), the density of water inside the cell is near identical to the density of fluid outside the cell whether the measurement is carried out in H_2_O or D_2_O-based solution (see (Delgado et al., 2013) for error estimations). Consequently, the latter term of *equation 1* approaches zero for both the H_2_O and the D_2_O-based buoyant mass measurements and the two consecutive measurements can be used to solve the first term of the equation, i.e. the dry mass and dry density of a cell.

### Measurement precision and stability

We characterized our measurement precision and stability using several approaches. First, we monitored live non-growing L1210 cells (mouse lymphocytic leukemia cells) by cooling our measurement system down to +4ºC to prevent growth and compositional changes. We measured the same cell repeatedly and defined our measurement error as the coefficient of variation during a 1h trace. With this approach our measurement errors in dry mass and dry density were ∼1.3% and ∼0.3%, respectively (Figure 1–figure supplements 1A&B). We did not observe measurement drifting even during a longer experiment (Figure 1–figure supplements 1A), suggesting a high level of measurement stability. Second, we monitored fixed cells at +37ºC and observed measurement errors that were close to those seen in live cells at +4ºC (Figure 1–figure supplement 1C). Third, we monitored the dry mass and density of 10 μm diameter polystyrene beads at +37ºC. This suggested similar dry mass measurement errors as the previous experiments with live and fixed cells, but lower errors in dry density measurements (Figure 1–figure supplement 1D). Thus, we estimate our dry mass measurement error to be ∼1.3%, which is comparable to current state-of-the-art QPM measurements (Liu et al., 2020; Reed et al., 2011). For dry density measurements, we estimate our measurement error to be ∼0.3%. We are not aware of any other single-cell methods capable of quantifying this biophysical feature of a cell. Furthermore, as we carry out dry composition measurements approximately every 75 s, we can further increase our precision by averaging multiple measurements while still capturing dry composition changes on timescales meaningful for studying dynamic and transient events such as mitosis.

### Measurement sensitivity to intracellular water exchange

In our typical measurement, a cell is exposed to D_2_O for 10-15 seconds. However, this time can vary, and if intracellular water exchange is not complete, the exposure time to D_2_O would influence our measurements. We tested this by correlating the measured dry mass and dry density to the time that a cell was exposed to the D_2_O-based media between each buoyant mass measurement. Whether examining end-point measurements of single L1210 cells in a population (Figure 1–figure supplement 2A), a single-cell trace of a live L1210 cell cooled to +4ºC (Figure 1–figure supplement 2B) or a single-cell trace of a L1210 cell at +37ºC (Figure 1–figure supplement 2C), the measured dry mass and dry density were independent of the time in D_2_O-based media. Thus, the rate of intracellular water exchange is not limiting our measurements, as also suggested by previous results in bacteria and yeast (Delgado et al., 2013).

## RESULTS

### Monitoring single cell’s dry mass and dry density

In order to continuously monitor the dry mass and dry density of single cells, we needed to periodically expose the cells to D_2_O, which can impact cell growth and viability. We first tested the sensitivity of L1210, BaF3 (mouse pro-B cell myeloma) and S-Hela (human adenocarcinoma) cells towards different concentrations of D_2_O by monitoring cell proliferation with live cell imaging. Under normal culture conditions, all three cell lines can tolerate continuous exposure to up to 30% D_2_O without changes to proliferation rate (Figure 2–figure supplement 1). Cells were still able to proliferate at higher D_2_O concentrations, albeit slowly.

**Figure 2.**
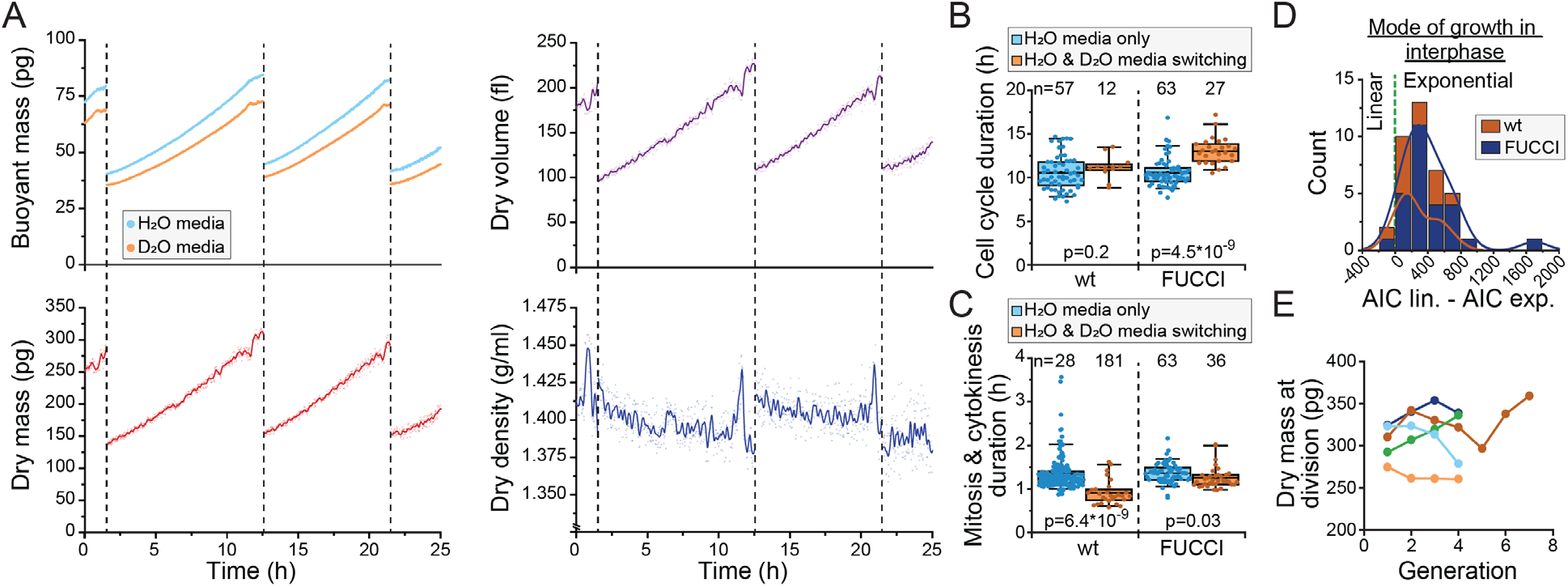
Monitoring buoyant mass, dry mass, dry volume and dry density of single cells. **(A)** An example dataset of an ancestral wt L1210 lineage tracked for its dry composition over two full cell cycles. Cell divisions are indicated with dashed vertical lines. Opaque points represent individual measurements and the solid lines represent smoothened data. **(B)** Cell cycle durations of wt and FUCCI L1210 cells grown in the SMR in normal media (blue) or with periodic D_2_O exposure (orange). Boxplot line: mean; box: interquartile range; whiskers: 5-95% range. p-values calculated using Welch’s t-test; n values refer to individual cells. **(C)** Same as (B), but data displays the combined duration of mitosis and cytokinesis. **(D)** Histogram of AIC of a linear fit – AIC of an exponential fit for wt and FUCCI L1210 cells in interphase (n=12 and 27 cells, respectively). Datapoints (individual cells) with positive values are better described by exponential rather than linear growth. **(E)** Dry mass at cell divisions across ancestral L1210 lineages. Each color displays a different experiment.

With the observed level of D_2_O tolerance, we hypothesized that we could monitor the dry density of a single cell growing inside the SMR as long as we minimize D_2_O exposure. To achieve this, we maintain the cell most of the time in the H_2_O-based media (Figure 1A), and our D_2_O-based media contains only 50% D_2_O. With this approach we were able to monitor the dry mass, dry volume and dry density of L1210 cells throughout multiple cell cycles (Figure 2A). We then examined if our measurement approach influences normal cell growth. Under these conditions, cell cycle durations of wild-type (wt) L1210 cells remained comparable to what we have observed previously in the SMR in the absence of D_2_O (Miettinen et al., 2019; Mu et al., 2020) (Figure 2B). However, when examining FUCCI (hGeminin-mKO) cell cycle reporter expressing L1210 cells, we observed slightly increased cell cycle durations. In addition, we observed that the duration of mitosis and cytokinesis was systematically shorter when cells were periodically exposed to D_2_O than what we have previously observed in other SMR experiments (Figure 2C).

Previous results examining mammalian cell growth in buoyant mass, dry mass or volume have all showed that cell growth in interphase is, on average, exponential (Cadart et al., 2018; Liu et al., 2020; Mu et al., 2020). For dry mass growth, this has only been examined using QPM. Utilizing our new dry mass growth measurements, we examined the mode of growth in wt and FUCCI L1210 cells in interphase. We compared linear and exponential growth models using Akaike information criterion (AIC) and found that exponential fits are a better model for our data in 37 out of 39 interphases (Figure 2D). We also examined the stability of cell size homeostasis in ancestral L1210 lineages with ≥3 full cell cycles and did not observe systematic changes in cell’s dry mass at division (Figure 2E). Together, these results show that we can monitor the dry mass and dry density of single cells over long time periods using our heavy water-based measurement approach, and while limited D_2_O exposure may influence cell proliferation rate, it does not radically alter cell size and growth regulation.

### Dry and buoyant mass correlate near-perfectly in interphase but not in mitosis

A common limitation of cell size and growth studies is the reliance on a single type of size measurement – for example, a cell’s dry mass or total volume. This can result in seemingly conflicting findings if cells change their composition. Our data enabled direct comparison of dry mass and buoyant mass measurements. We first examined the correlation between dry mass and buoyant mass on population level by carrying out end-point measurements of single cells. We observed a high degree of correlation (R^2^>0.97) for L1210, BaF3 and S-Hela cell populations (Figure 2–figure supplement 2A). We then correlated dry mass and buoyant mass within our L1210 single-cell growth traces and observed a very high degree of correlation in interphase of each cell (R^2^∼0.99), but lower and more variable correlations in mitosis and cytokinesis (R^2^∼0.5-0.9) (Figure 2–figure supplements 2B&C). Together, these observations indicate that buoyant mass is an accurate proxy for cell’s dry mass in interphase, but not necessarily in mitosis.

Cells’ dry mass and buoyant mass will correlate if dry density does not change, assuming that measurements are carried out in normal cell culture media or other water-based solution with density close to water (see equation 1). Indeed, in our population level data, dry density displayed little variability, with coefficient of variability around 1.6% for L1210, BaF3 and S-Hela cells (Figure 2–figure supplement 2D). This is approximately 15-fold less variability than observed for dry mass. The low variability in dry density within a proliferating cell population represents the tight regulation of cells’ molecular composition.

### Cells transiently lose dry mass and increases dry density in early mitosis

We then examined dry composition behavior(s) in mitosis. We detected mitotic stages based on acoustic scattering from the cell, which reports approximate mitotic entry and the metaphase-anaphase transition (Kang et al., 2019) (Figure 3–figure supplement 1). We also validated the metaphase-anaphase transition timing by examining the degradation of the hGeminin-mKO cell cycle reporter in the FUCCI L1210 cells (Figure 3–figure supplement 1). Curiously, we observed that many, but not all, L1210 cells lost dry mass and, especially, dry volume following mitotic entry, resulting in increased dry density in early mitosis (Figures 3A-C; Figure 3-source data 1). This was observed in both wild-type and FUCCI L1210 cells, although the FUCCI cells displayed larger dry composition changes in mitosis (Figure 3B). On average, the FUCCI L1210 cells lost ∼4% of dry mass and increased dry density by ∼2.5%, and these changes took place in approximately 15 minutes (Figure 3C). In extreme cases, cells lost ∼8% of their dry mass while increasing dry density by ∼4%. These changes were transient, as cells recovered their lost dry mass and decreased their dry density at metaphase-anaphase transition (Figure 3B). The recovery of dry mass at metaphase-anaphase transition was faster than the dry mass growth observed prior to mitosis, suggesting that this ‘recovery growth’ is distinct from normal cell growth. After the metaphase-anaphase transition, dry mass growth continued during cytokinesis. The total amount of dry mass accumulated in mitosis and cytokinesis was large, over 10% of the dry mass accumulated in the whole cell cycle (Figure 3D). The extent of mitotic dry mass loss in early mitosis was coupled to the extent of corresponding dry density increase (Figure 3E). Importantly, the increase in dry density uncoupled buoyant mass and dry mass measurements (equation 1), causing buoyant mass to increase in early mitosis even in the cells which lost dry mass (Figure 3–figure supplements 2A-C).

**Figure 3.**
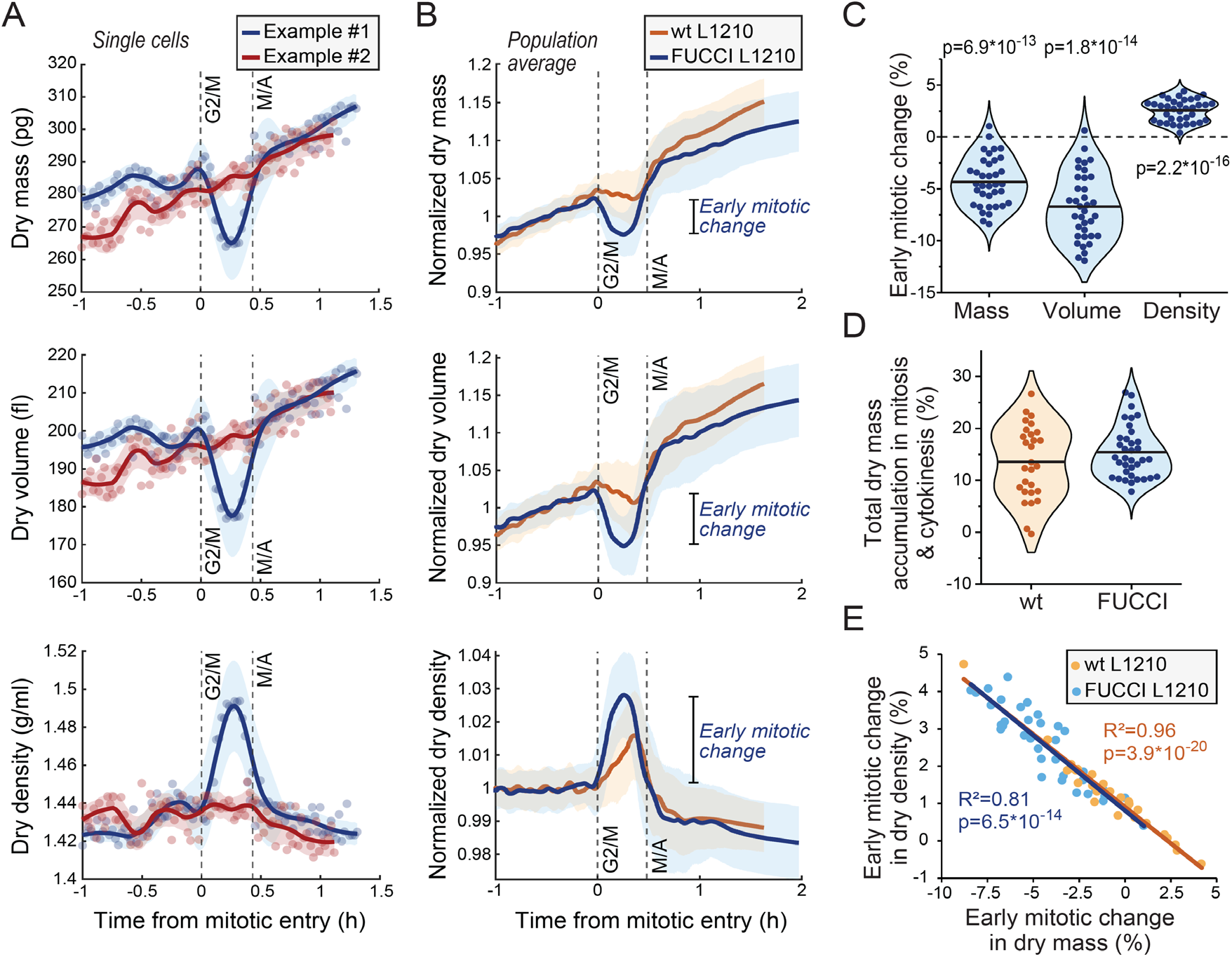
Cells transiently lose dry mass and increase dry density in early mitosis. **(A)** Two example FUCCI L1210 cells with different dry mass (top), dry volume (middle) and dry density (bottom) behaviors in early mitosis. Opaque points represent individual measurements; thick line and shaded area represent smoothened data (mean±SD). Dashed vertical lines indicate approximate G2/M and metaphase-anaphase (M/A) transitions. **(B)** Population average dry composition behavior in wt and FUCCI L1210 cells. Thick line and shaded area represent mean±SD. N=31 cells from 19 independent experiments for the wt cells; N=36 cells from 13 independent experiments for the FUCCI cells. **(C)** Quantifications of dry composition changes in early mitosis in FUCCI L1210 cells. Dots represent individual cells; horizontal line represents mean; data is same as in panel (B); p-values calculated using one sample t-test and represent difference from zero. **(D)** Dry mass accumulation from mitotic entry to cell division relative to the dry mass accumulated in the whole cell cycle. Dots represent individual cells; horizontal line represents mean; data is same as in panel (B). **(E)** Correlation between dry mass and dry density change in early mitosis for wt and FUCCI L1210 cells. Data is same as in panel (B); correlation p-values calculated using ANOVA. Raw data can be found in Figure 3—source data 1.

### Mitotic dry composition changes display large, non-genetic cell-to-cell variability

The early mitotic dry composition changes were not observed uniformly across all wt and FUCCI L1210 cells. Approximately 7% of all L1210 cells measured (10 out of 67 cells) displayed little to no dry density increase in early mitosis, and the cell-to-cell variability in mitotic dry composition changes was large (Figure 3C). We also observed mother-daughter pairs where the mother cell displayed mitotic changes in its dry composition but the daughter cell did not, or *vice versa* (Figure 3–figure supplements 2A-C). The difference in mitotic dry composition behavior between individual cells did not correlate with experiment-specific noise levels (Figure 3–figure supplement 2D). Together, these results indicate a large degree of non-genetic cell-to-cell variability in the regulation of mitotic dry composition. Furthermore, since the dry composition changes were not present in all cells, they are not required for a successful cell division.

We also monitored the dry composition of BaF3 and THP-1 (human acute monocytic leukemia) cells through mitosis. The BaF3 cells displayed a large degree of cell-to-cell variability in mitotic dry composition behavior. While some individual BaF3 cells lost ∼4% of their dry mass during early mitosis, on average the BaF3 cells did not lose dry mass in mitosis but growth rather stopped during early mitosis, which was followed by a rapid recovery at metaphase-anaphase transition (Figure 3–figure supplement 3A-E). In THP-1 cells, on average, cells lost ∼5% of their dry mass, which was also recovered at metaphase-anaphase transition (Figure 3–figure supplement 3F-J). In both cell types, dry density transiently increased in early mitosis. Thus, the mitotic loss of dry mass and increase in dry density are not specific to L1210 cells. More broadly, our dry mass measurements suggest that early mitosis is coupled to previously unrecognized secretion of cellular components.

### Mitotic dry composition changes do not require morphological changes

We investigated the mechanistic basis for the mitotic changes in dry composition using L1210 cells as our model system. We first examined the influence of mitotic progression and mitotic morphological changes to cell’s dry composition. We arrested wt L1210 cells to prometaphase using 5 μM kinesin inhibitor S-trityl-l-cysteine (STLC) and validated mitotic arrest by quantifying the fraction of mitotic cells using DNA and p-Histone H3 (Ser10) labeling (Figure 4A). When monitoring the dry composition of STLC treated cells, we observed slightly larger, but statistically not significantly different, loss of dry mass and increase in dry density upon mitotic entry as in control cells (Figures 4B&C). However, the mitotically arrested cells did not recover their dry composition, and following the initial changes, the STLC treated cells displayed slow dry mass accumulation and steady dry density for the rest of the mitotic arrest. These results verify that the increase in dry density and decrease in dry mass take place before the completion of prometaphase, and that mitotic progression is required for the rapid recovery of dry mass and dry density.

**Figure 4.**
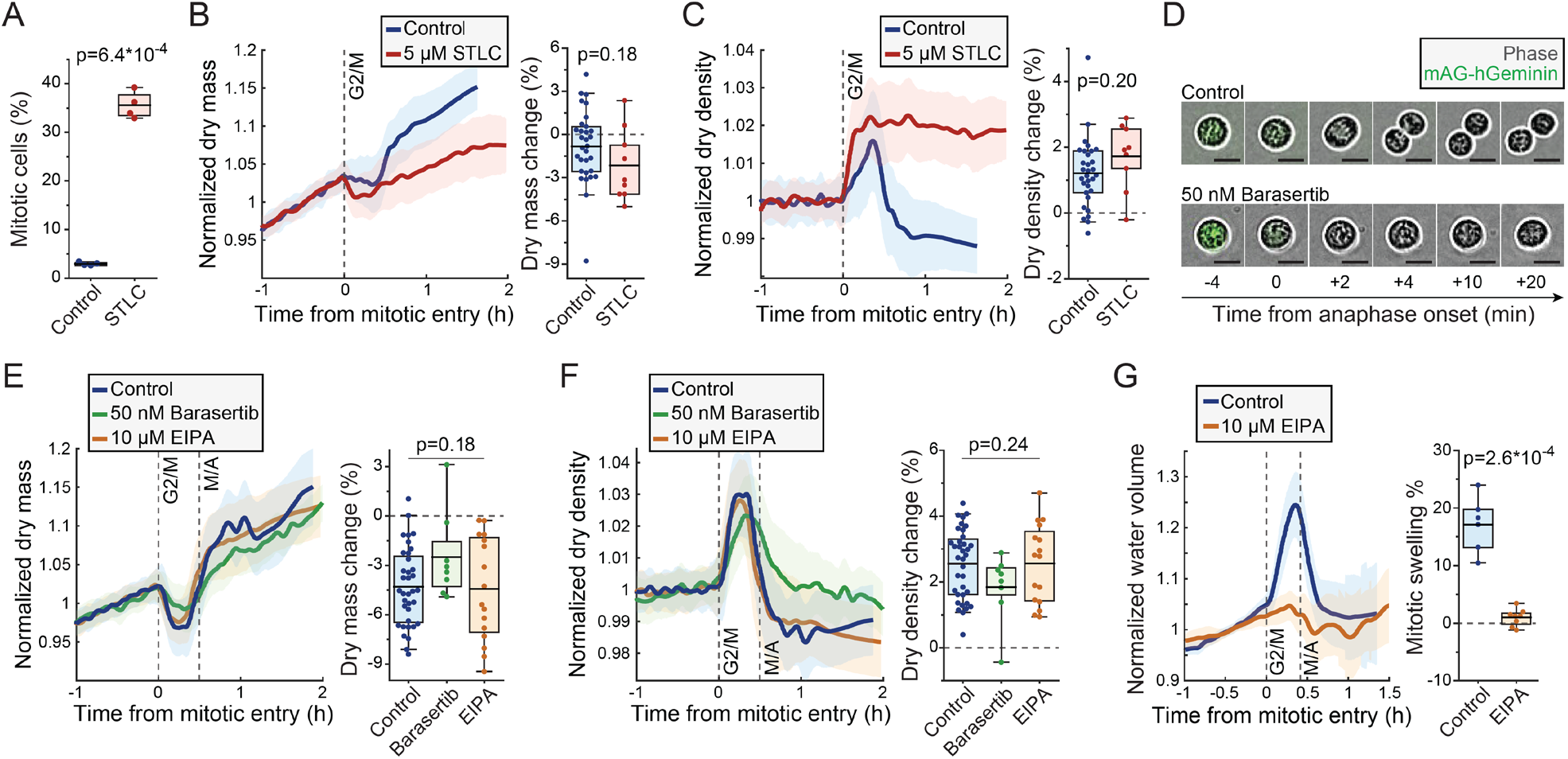
Mitotic dry mass loss and dry density increase are independent of morphological changes. **(A)** % of mitotic cells in a wt L1210 population following 5 h control or STLC treatment. N=4 independent cultures; p-value calculated using Welch’s t-test. **(B)** Normalized dry mass behavior for control and STLC treated wt L1210 cells (left) and quantifications of early mitotic dry mass changes (right). N=31 cells from 19 independent experiments for control; N=9 cells from 9 independent experiments for STLC; p-value calculated using Welch’s t-test. **(C)** Same as panel (B), but for dry density. **(D)** Representative phase contrast and mAG-Geminin reporter images of control and Barasertib treated FUCCI L1210 cells in mitosis. N>20 cells from 3 independent experiments. **(E)** Normalized dry mass behavior for FUCCI L1210 cells treated with indicated chemicals (left) and quantifications of early mitotic dry mass changes (right). N=36 cells from 13 independent experiments for control; N=8 cells from 8 independent experiments for Barasertib; N=16 cells from 15 independent experiments for EIPA; p-value calculated using ANOVA. **(F)** Same as panel (E), but for dry density. **(G)** Normalized intracellular water volume behavior for control and EIPA treated FUCCI L1210 cells (left) and quantifications of mitotic cell swelling (right). N=6 cells from 5 independent experiments for control; N=8 cells from 8 independent experiments for EIPA; p-value calculated using Welch’s t-test. In dry mass, dry density and water volume traces, the thick line and shaded area represent mean±SD; Boxplot line: mean; box: interquartile range; whiskers: 5-95% range. Raw data can be found in Figure 3—source data 1.

As the recovery of dry composition required mitotic progression, we asked if the recovery is coupled to cell elongation. To this end, we inhibited cytokinesis in FUCCI L1210 cells using 50 nM Aurora B inhibitor Barasertib. We validated the inhibition of cytokinesis by imaging cell morphology as cells proceeded through metaphase-anaphase transition, as indicated by the degradation of the hGeminin-mKO signal (Figure 4D). When monitoring cells’ dry composition, the Barasertib-treated cells displayed slightly lower, but statistically not significantly different, loss of dry mass and increase in dry density as untreated FUCCI L1210 cells (Figures 4E&F). Although the duration of mitosis was lengthened in some Barasertib-treated cells, resulting in slower recovery of dry mass and dry density, the dry composition changes that took place in early mitosis eventually recovered to similar levels as seen in untreated cells. Thus, the mitotic changes in cell’s dry composition cannot be explained by cell elongation, although the rate of dry composition recovery in anaphase may be influenced by cell elongation or Aurora B activity.

Another major morphological change that mitotic cells undergo is the mitotic cell swelling, where cells take up water and can increase their total volume by 10-20% (Son et al., 2015; Zlotek-Zlotkiewicz et al., 2015). This takes place simultaneously with the dry composition changes, i.e. cells swell from prophase to metaphase and recover in anaphase. To investigate if mitotic cell swelling is required for the dry composition changes, we inhibited mitotic swelling using 10 μM sodium-hydrogen exchanger inhibitor 5-(N-ethyl-N-isopropyl)-Amiloride (EIPA). We first validated that EIPA prevents mitotic cell swelling by monitoring approximate intracellular water volume (Son et al., 2015). This revealed that the FUCCI L1210 cells swell significantly in early mitosis, but this is near completely prevented by EIPA treatment (Figure 4G). However, when we monitored the dry mass and density of FUCCI L1210 cells in the presence of EIPA, we did not observe any changes in the amount of dry mass lost or dry density increased in early mitosis when compared to control cells (Figures 4E&F). Therefore, the mitotic dry composition changes are not due to mitotic cell swelling. Finally, we validated that the mitotic perturbations used here did not influence the typical dry density of the cells (Figure 4–figure supplement 1).

### Lysosomal exocytosis is increased in early mitosis

Cells undergo a constant influx and efflux of dry mass through the flux of ions, metabolites, proteins and other components that are taken up and secreted by a cell. When evaluating potential mechanisms for the mitotic loss of dry mass, we first considered the extent of dry mass loss in mitosis, which was ∼8% in extreme cases. Macromolecules would be the most likely form of mass lost, as they constitute most of cell’s dry mass. In contrast, inorganic ions are an unlikely form of dry mass lost, as these constitute only ∼1% of cell’s dry mass (Cooper, 2013). Cells could secrete metabolites or small molecules in mitosis, but at least the secretion of lactate, a key metabolite secreted by cells, is not increased in mitotic L1210 cells (Kang et al., 2020). Furthermore, it seems unlikely that cells could lose large amounts of ions and/or small molecules while increasing osmotic pressure, which is required for the mitotic cell swelling. We therefore focused our study on mechanisms that secrete macromolecules.

Macromolecule secretion is largely due to exocytosis, the process where intracellular vesicles or organelles merge with the plasma membrane to release their contents to the extracellular space. This is balanced by the reverse process called endocytosis. Most studies have found that endocytosis stops in mitosis, more specifically from mitotic entry until anaphase (Fielding et al., 2012; Schweitzer et al., 2005; Warren et al., 1984). This corresponds to the timing of dry mass loss that we have observed. However, contradictory results on mitotic endocytosis have also been reported (Boucrot and Kirchhausen, 2007). Changes in exocytosis have not been studied in similar detail during mitosis, but lysosomal exocytosis, where cells secrete lysosomal contents by fusing lysosomes with the plasma membrane (Reddy et al., 2001), has been suggested to increase in late mitosis (Nugues et al., 2018). Notably, lysosomal exocytosis has also been shown to secrete fatty acids (which are low in dry density) out of the cell (Cui et al., 2021), which could increase the dry density of a cell.

We hypothesized that the loss of dry mass in early mitosis is driven by increased exocytosis of components such as lysosomes. Lysosomal exocytosis, and the fusion of lysosomes with the plasma membrane, can be quantified by monitoring the presence of the lysosomal membrane protein LAMP-1 on cell surface (Reddy et al., 2001; Samie and Xu, 2014). We immunolabelled LAMP-1 in freely proliferating, live L1210 cells and validated the surface-specific labeling using microscopy (Figure 5A; Figure 5–figure supplement 1). Quantifications of the microscopy indicated that mitotic cells had more LAMP-1 on their surface than interphase cells (Figure 5B). Furthermore, the amount of LAMP-1 on cell surface increased from early mitosis (prophase, prometaphase and metaphase) to late mitosis (anaphase) (Figure 5B), indicating that cell surface LAMP-1 accumulates throughout early mitosis. This is in contrast to the previous report suggesting that lysosomal exocytosis increases specifically in late mitosis (Nugues et al., 2018).

**Figure 5.**
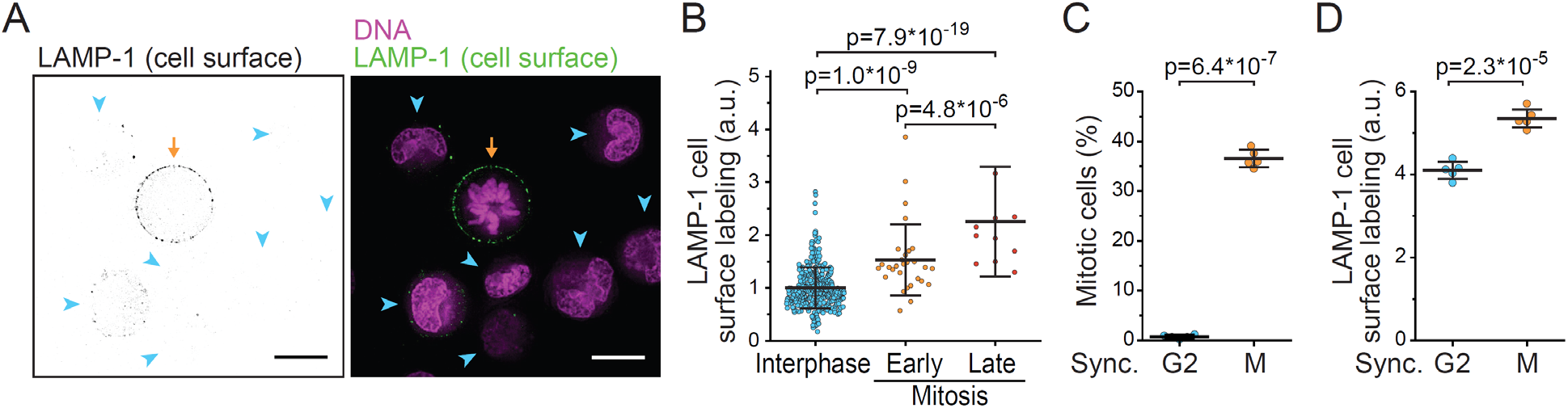
Lysosomal exocytosis is increased in early mitosis. **(A)** A representative image of surface LAMP-1 immunolabeling alone (left) and together with DNA labeling (right) in live L1210 cells. Orange arrow indicates a mitotic cell, blue arrowheads indicate interphase cells. Scalebars depict 10 μm. **(B)** Microscopy quantifications of cell surface LAMP-1 immunolabeling in unsynchronized L1210 cells. Early mitosis refers to prophase, prometaphase and metaphase; late mitosis refers to anaphase. N=380, 29 and 11 cells for interphase, early and late mitotic cells, respectively; data pooled from 2 independent experiments; p-values calculated using ANOVA followed by Sidakholm posthoc test. **(C)** % of mitotic cells in L1210 cell populations following synchronization to G2 or early mitosis. N=5 independent cultures; p-value calculated using Welch’s t-test. **(D)** Flow cytometry quantifications of L1210 cell population surface LAMP-1 immunolabeling following synchronization to G2 or early mitosis. N=5 independent cultures; p-value calculated using Welch’s t-test. In all figures, line and whiskers indicate mean±SD.

To further support the observation that lysosomal exocytosis is increased in early mitosis, we synchronized L1210 cells to G2 and early mitosis using RO-3306 (CDK1 inhibitor) and STLC, respectively (Figure 5C). Notably, STLC arrests cells in prometaphase, where the mitotic dry composition changes are taking place. We then immunolabelled cell surface LAMP-1 and quantified the labeling using flow cytometry. Cells arrested to early mitosis displayed higher levels of LAMP-1 on their surface than cells arrested to G2 (Figure 5D). We also examined cell surface LAMP-1 immunolabeling in BaF3 and THP-1 cells after cell cycle synchronizations to G2 or early mitosis. In both cell lines, LAMP-1 labeling was higher in early mitosis than in G2 (Figure 5–figure supplement 2). Thus, lysosome exocytosis is increased in early mitosis.

### Chemical inhibition of lysosomal exocytosis prevents mitotic dry composition changes

We then investigated if inhibition of lysosomal exocytosis prevents the dry composition changes taking place in mitosis. For this, we utilized 10 μM Brefeldin A, an inhibitor of vesicle-mediated protein trafficking and exocytosis (Hendricks et al., 1992), and 15 μM Vacuolin-1, a small molecule inhibitor of lysosomal exocytosis (Cerny et al., 2004). The ability of Vacuolin-1 to preventing lysosomal exocytosis has been controversial (Cerny et al., 2004; Cui et al., 2021; Huynh and Andrews, 2005), and we first validated that lysosomal exocytosis is inhibited by Brefeldin A and Vacuolin-1. Both chemicals prevented the accumulation of LAMP-1 on plasma membrane in L1210 cells (Figure 6A; Figure 5–figure supplement 1), indicating that lysosomal exocytosis was inhibited.

**Figure 6.**
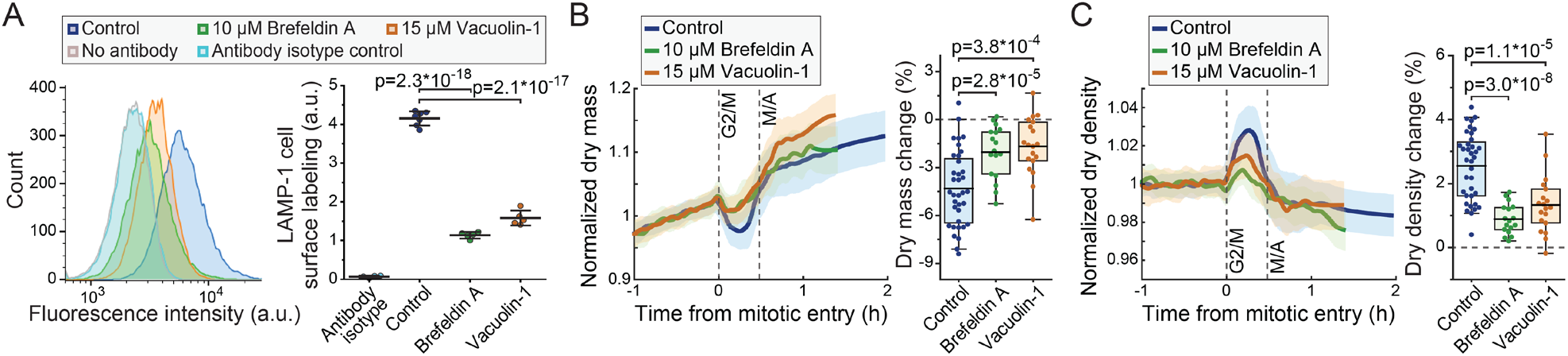
Inhibitors of lysosomal exocytosis prevent mitotic dry composition changes. **(A)** Representative histograms (left) and quantifications (right) of live L1210 cell surface LAMP-1 immunolabeling following indicated, 4 h long chemical treatments. Line and whiskers indicate mean±SD; N=4-7 independent cultures. **(B)** Normalized dry mass behavior for FUCCI L1210 cells treated with indicated chemicals (left) and quantifications of early mitotic dry mass changes (right). N=36 cells from 13 independent experiments for control; N=17 cells from 17 independent experiments for Brefeldin A; N=19 cells from 19 independent experiments for Vacuolin-1; p-values calculated using ANOVA followed by Sidakholm posthoc test; Thick line and shaded area represent mean±SD; Boxplot line: mean; box: interquartile range; whiskers: 5-95% range. **(C)** Same as panel (B), but for dry density. Raw data can be found in Figure 3—source data 1.

We then monitored the dry composition of FUCCI L1210 cells in the presence of Brefeldin A or Vacuolin-1. Both chemicals prevented the mitotic changes to cells’ dry composition (Figure 6B&C) without preventing cell divisions. The chemicals did not cause clear changes to the baseline dry density of cells (Figure 6–figure supplement 1). Thus, these results indicate that lysosomal exocytosis is, at least partly, responsible for the loss of dry mass and increased dry density in early mitosis. However, we note that other forms of exocytosis may also be involved in the dry composition changes.

## DISCUSSION

Here we have developed a method using the SMR to monitor the dry mass, dry volume and dry density of suspension grown cells over long periods of time with high precision. To the best of our knowledge, this is the first method capable of non-invasively measuring cell’s dry volume or the density of cell’s dry mass. We anticipate several uses for this method. First, our dry density measurements provide an orthogonal approach for studying the macromolecular composition of cells. While dry density measurements lack molecular specificity, they are non-invasive, quantitative, and have high temporal resolution. Together, these qualities make our measurements valuable for studying the dynamics of molecular composition changes, especially in situations where rapid changes cannot be easily resolved by more typical population level measurements, such as mass spectrometry. Second, our method provides an additional approach for measuring dry mass. QPM has several advantages over our method, including higher throughput and suitability for adherent cells, but our method is comparable in precision and has high temporal resolution without phototoxicity. This enables us to observe dry mass dynamics that may be difficult to capture with other methods. Third, the ability of our method to measure cellular dry volume could enable precise, specific and non-invasive quantifications of single-cell water content, provided it is coupled to a measurement of total cell volume.

Using our new method, we have expanded our previous study of mammalian cell growth dynamics in mitosis. Importantly, the mitotic dry mass dynamics differ from our previously reported buoyant mass dynamics due to a transient change in dry density in early mitosis (equation 1). This finding reconciles our previous mitotic buoyant mass measurements (Miettinen et al., 2019) with the QPM-based dry mass measurements by others (Liu et al., 2020; Zlotek-Zlotkiewicz et al., 2015). The overall dry mass growth remains high in mitosis and cytokinesis, and for L1210 cells over 10 % of the total dry mass growth in the cell cycle takes place in mitosis and cytokinesis. This supports our previous main conclusion that average cell growth rates in mitosis and cytokinesis are comparable to those observed in interphase, which is consistent with the high protein synthesis rates reported for unperturbed mitosis (Coldwell et al., 2013; Miettinen et al., 2019; Stonyte et al., 2018; Sun et al., 2019). In interphase, dry density is conserved, making buoyant mass an excellent proxy for dry mass.

A key discovery made here is the significant loss of dry mass in early mitosis, which can be up to 8% of cell’s total dry mass. Importantly, a loss of dry mass in early mitosis has been reported in adherent mammalian cells studied using QPM (Liu et al., 2020), but the higher temporal resolution of our approach enables us to examine this phenotype in more detail. Our results reveal a high degree of cell-to-cell variability in the mitotic loss of dry mass and cells can divide without losing mass. Mechanistically, we show that exocytosis of lysosomes is increased in early mitosis and the loss of dry mass is dependent on exocytosis. While lysosomal exocytosis is at least partly responsible for the loss of dry mass, other mechanisms, especially other forms of exocytosis, may be involved. In addition, the loss of dry mass in early mitosis could be partly explained by decreased endocytosis, as the total dry mass of a cell is defined by the balance between uptake and secretion processes, such as endo- and exocytosis, respectively. Our results also show that cells are able to recover dry mass at a high rate at metaphase-anaphase transition. Interestingly, the exocytosis of lysosomes is typically followed by endocytosis, which removes lysosomal membranes from plasma membrane (Idone et al., 2008; Tam et al., 2010). Such a compensatory endocytosis could explain the rapid recovery of dry mass at metaphase-anaphase transition, but further quantitative studies on endo- and exocytosis are needed to confirm this.

Why would cells exocytose lysosomal contents and a significant amount of total cellular biomass in mitosis? We hypothesize three potential reasons for this: 1) Lysosomal exocytosis acts in plasma membrane repair (Reddy et al., 2001), and the lysosomal exocytosis could enable a temporarily increase in plasma membrane area for mitotic cell swelling. 2) Lysosomal exocytosis is used for secretion of various proteins and enzymes in to the extracellular space (Blott and Griffiths, 2002), and mitotic cells may secrete extracellular matrix modifying enzymes in order to ensure an appropriate physical milieu for cell division. 3) Lysosomal exocytosis has been shown to clear the cell of excessive/toxic cellular proteins, such as α-synuclein (Tsunemi et al., 2019) and tau (Xu et al., 2020), and mitotic lysosomal exocytosis may act as a ‘reset’ that enables the daughter cells to be born with a minimal load of useless/harmful contents. Consequently, non-proliferating cells, such as neurons, could be more prone to suffer from excessive/toxic protein buildup, and the mechanism(s) that promotes mitotic lysosomal exocytosis could act as a candidate drug target for promoting cellular clearance of toxic proteins. Notably, lysosomal exocytosis is also linked to a variety of lysosomal and lipid storage disorders (reviewed in (Samie and Xu, 2014) and (Settembre and Ballabio, 2014)), and understanding the mechanism(s) of mitotic lysosomal exocytosis could have implications for these diseases as well.

## MATERIALS AND METHODS

### Key resources table

The key resources table is attached to this manuscript as an excel file.

### Cell culture and media composition

L1210, BaF3, THP-1 and S-Hela cells were cultured in RPMI media (Thermo Fisher Scientific, #11835030) supplemented with 10% FBS, 1 mM sodium pyruvate, 10 mM HEPES and antibiotic/antimycotic. This served as the H_2_O-based media for experiments and the D_2_O-based media was made to an identical composition using RPMI media powder (Thermo Fisher Scientific, #31800022) and heavy water (Sigma-Aldrich, #151882). D_2_O-based media pH was adjusted to 7.4 with HCl and NaOH, and the media was filtered with 0.2 μm filter. To minimize cell exposure to heavy water in the SMR experiments, the D_2_O-based media was diluted with the normal media to contain 50% heavy water. All experiments were started using cells growing exponential at a confluency of 250.000-500.000 cells/ml.

### Cell confluency and morphology imaging

Cell population growth in medias with increasing amounts of D_2_O was measured by imaging cell confluency on a plate over ∼3 days. Confluency was monitored using the IncuCyte live cell analysis imaging system by Sartorius. Imaging was carried out every 3 hours using 4X objective and confluency was analyzed using IncuCyte S3 2017 software’s standard settings. The confluency data were then used to calculate confluency doubling time, which was used as a proxy for cell cycle duration. For validations of cell morphology in mitosis, cells were imaged every 2 minutes with 20X objective using the IncuCyte. Metaphase-anaphase transition was detected by simultaneous imaging of the FUCCI cell cycle sensor on FITC channel. The example images were overlaid and processed in the IncuCyte S3 2017 software.

### Chemical perturbations, cell cycle synchronizations, cell cycle analyses

Chemical treatment concentrations are indicated in main text and supplier details can be found in the key resources table. For SMR experiments, the cells were not pretreated with the chemical inhibitors, and all treatments started at the beginning of each SMR experiment. The treatment time until mitosis therefore varies between experiments, with typical range being 2-4 hours. Apart from EIPA treatments, all SMR experiments with chemical inhibitors lasted only for one cell division.

For cell cycle synchronization in G2 and mitosis, cells growing at ∼400.000 cells/ml confluency were first arrested in G2 using 5 μM RO-3306 treatment. For L1210 and BaF3 cells, this treatment lasted for 6 h; for THP-1 cells this treatment lasted for 20 h. The cells were then washed twice with cold PBS and moved in to fresh media containing either 5 μM RO-3306 (for G2 arrest) or 5 μM STLC (for prometaphase arrest). For L1210 and BaF3 cells, this treatment lasted for 1 h; for THP-1 cells this treatment lasted for 2 h. After this, the cells were either fixed for cell cycle analysis or immunolabelled with anti-LAMP-1 antibody in drug containing media. Cell cycle arrest in G2 and mitosis was validated using flow cytometry based detection of DNA and phospho-Histone H3 (Ser10) labeling, as detailed before (Miettinen et al., 2019). Results in Figure 4A, Figure 5C and Figure 5–figure supplement 2A&C represent the fraction of cells that were p-Histone H3 (Ser10) positive and contained 4N DNA content, as evaluated based on a comparison to freely proliferating cells.

### Measurements of cell surface localized LAMP-1

In L1210 and BaF3 cells, the cell surface localized LAMP-1 protein was detected using immunolabeling with rat monoclonal anti-LAMP-1 antibody conjugated to Alexa Fluor 488 (Thermo Fisher Scientific; RRID:AB_657536). The cells were moved to fresh media (containing chemical treatments, where applicable) and incubated with the antibody at 1:50 dilution for 40 min on ice, after which the cells were washed twice with fresh media and either imaged or measured using flow cytometry. For antibody isotype control, we used rat monoclonal IgG2a kappa Isotype Control antibody conjugated to Alexa Fluor 488 (Thermo Fisher Scientific; RRID:AB_493963). For THP-1 cells, the immunolabeling protocol was otherwise identical, but the immunolabeling was carried out with mouse monoclonal anti-LAMP-1 antibody conjugated to Alexa Fluor 488 (Thermo Fisher Scientific; RRID:AB_2016657) or with corresponding isotype control (RRID:AB_470230).

When imaging the LAMP-1 immunolabelled cells, the last 20 min of antibody labeling was carried out together with DNA labeling using NUCLEAR-ID red DNA stain at 1:250 dilution (Enzo Life Sciences, Cat#ENZ-52406). For microscopy, we used DeltaVision wide-field deconvolution microscope with standard filters (FITC, APC), 100× objective and an immersion oil with a refractive index of 1.516. For each image, we collected 15 z-slices with 0.2 μm spacing, deconvolved the images using SoftWoRx 7.0.0 software and selected a middle z-slice for visualization. For flow cytometry-based detection of LAMP-1, DNA was not stained and LAMP-1 immunolabeling was detected using BD Biosciences flow cytometer LSR II HTS with excitation laser at 488 nm and emission filter at 530/30.

Image quantifications were carried out manually from single z-layers by cropping out each cell and calculating the average LAMP-1 labeling intensity per cell in ImageJ (version 2.0.0-rc-69/1.52p). z-layers with intracellular labeling were rare and not used for analysis. Cells were classified as interphase, early or late mitotic cells based on their DNA morphology. After this, the background signal was removed from the cell signals, and the LAMP-1 labeling of each cell was normalized to the average intensity observed in all interphase cells within the same image (typical image contained ∼19 interphase cells).

### SMR operation for dry mass and dry density measurements

Basic SMR build and operation was identical to that reported before (Kang et al., 2020; Kang et al., 2019; Miettinen et al., 2019). The concept of dry mass and dry density measurements is detailed in (Delgado et al., 2013). For dry mass and dry density measurements, the bypass channel on one side of the SMR cantilever is filled with H_2_O-based media and the other side with D_2_O-based media. For end-point single-cell measurements, the cells are first loaded on the H_2_O-based media side of the device at ∼300.000 cells/ml concentration. A cell is then flown through the SMR to obtain a measurement in the H_2_O-based media. On the other side of the cantilever the cell is diluted in D_2_O-based media, which is in excess to the H_2_O-based media to maximize dilution. In a typical end-point measurement, the cell spends 5-8 seconds in the D_2_O-based media. The cell is then flown back through the SMR to obtain a second measurement, this time in the D_2_O-based media. After this, both sides of the cantilever are flushed with fresh media & cell(s) and the cycle is repeated. For single-cell monitoring, the protocol is the same as for end-point measurements, except that the cell is maintained for ∼55 s in the H_2_O-based media and ∼15 s in D_2_O-based media. Time in D_2_O-based media is kept longer than in end-point measurements, as this increases the long-term stability of single-cell trapping. Importantly, the SMR baseline frequency is recorded also prior and after every cell measurement. As the SMR baseline frequency is directly proportional to media density, this allows us to adjust each measurement for local mixing of H_2_O and D_2_O-based medias around the cell.

### Calibration and calculation of dry mass and dry density from SMR data

The two buoyant mass measurement (SMR frequency response to a cell flowing through the cantilever) and media density measurements (SMR frequency baseline immediately before and after the cell flows through the cantilever) were used to calculate dry mass, dry volume and dry density according to *equation 1*, assuming that extracellular fluid (culture media) density is equal to the density of water (H_2_O or D_2_O) inside the cell (Delgado et al., 2013). In order to calibrate the SMR baseline, sodium chloride solutions of known density were measured at RT and used to generate a calibration curve for SMR baseline. In addition, the buoyant mass measurements were calibrated using 10 μm diameter monodisperse polystyrene beads (Thermo Fisher Scientific, Duke Standards), where the volume and density of the measured particle are known (Mu et al., 2020).

### Dry mass and dry density measurement resolution estimations

The precision of dry mass and dry density measurements was evaluated by repeatedly measuring the same, non-growing particle. As there are many sources of noise, three types of non-growing particles were used: a fixed L1210 cell measured at +37°C (fixation in 4% PFA in PBS for 10 min followed by a wash with PBS), a live L1210 cell measured at +4°C, and a 10 μm diameter polystyrene bead measured at +37°C. Each particle was tracked for dry mass and dry density similarly to that described above for live cells. For each experiment, 1h long section of the trace was used to calculate the measurement precision, presented as the coefficient of variability. When evaluating experiment specific measurement noise (Figure 3–figure supplement 2D), we fitted the last 1h of dry mass data in interphase with a linear fit and calculated the SD of the residuals. Note that this calculation includes both technical and biological noise.

### Intracellular water volume approximation using the SMR

Approximate intracellular water volume of FUCCI L1210 cells was measured in the SMR by growing the cells in culture media containing 40% OptiPrep Density Gradient Medium (Sigma-Aldrich). The OptiPrep containing media has a solution density which is closer to cell’s dry density than the density of water, thus making the SMR-based buoyant mass measurement predominantly sensitive towards intracellular water volume (*equation 1*) (Son et al., 2015). Apart from the media change, the SMR was operated as before (Miettinen et al., 2019), and the measurements were calibrated using 10 μm diameter monodisperse polystyrene beads (Thermo Fisher Scientific, Duke Standards).

### Data analysis, smoothing & plotting

Each cell trace analysis was limited to data starting from 1 h before mitotic entry and ending at 2 h after mitotic entry or at cell division. The very last datapoint prior to abscission was excluded in order to avoid analysis of cells during the final abscission. The traces were then normalized to their average value observed in the 1 h period before mitotic entry, and the traces were overlayed. When plotting the dry composition mean±SD values for each control and/or treatment condition, the moving and locally weighted average was plotted together with identical length moving standard deviation. For quantifications of dry composition changes in early mitosis, we identified the time point in mitosis where, on average, each control/treatment reached the minimum dry mass. The average of 5 datapoints at the maximal effect size was then compared the average of 5 datapoints immediately prior to mitotic entry (inclusive of the indexed mitotic entry timepoint). The difference between these two values is defined as ‘early mitotic change’. Note that the maximal effect size was always achieved prior to metaphase-anaphase transition. Same smoothing and quantification approach was also used for processing the intracellular water volume measurements. For single-cell examples, the cut-out traces were smoothened using LOESS smoothing.

The mode of cell growth was analyzes by comparing an exponential and a linear fit to the raw dry mass data in interphases. Akaike information criterion was used to define which fit is better (Liu et al., 2020). Only cells with full cell cycle (from birth to division) were used. All data processing and smoothing was done using MATLAB (versions 2017b and 2020b), and figures were plotted using MATLAB and Origin (version 2021b).

### Statistics

The statistical tests used, the results of these tests and detailed information about replicate numbers are indicated in each figure and figure legend. All statistical tests were carried out using Origin (version 2021b) software. Appropriate sample sizes were not computed before experiments and sample sizes were defined by practical considerations (e.g. experimental time requirements). We did not blind-control experiments, as most work was carried out by a single individual. When trapping a cell in the SMR for dry mass and dry density measurements, we trap randomly the first normal-sized cell that flows through the SMR after cells are flushed in to the system.

### Source data

All dry mass and dry density traces of live cells, including those where cells are treated with chemical inhibitors, are available in Figure 3-source data 1.

## Data availability

All data are included in the manuscript and supporting source data files.

## Acknowledgements

We thank Dr. Ye Zhang for assistance with data analysis. S.R.M. received funding and support from the MIT Center for Cancer Precision Medicine, Virginia and D.K. Ludwig Fund for Cancer Research, Cancer Systems Biology Consortium (CA217377) and Cancer Center Support (core) Grant P30-CA14051 from the National Cancer Institute.

## Author contributions

Teemu P Miettinen: Conceptualization, Data curation, Formal analysis, Supervision, Validation, Investigation, Visualization, Methodology, Writing—original draft, Writing—review and editing. Kevin Ly: Data curation, Formal analysis, Visualization, Writing—review and editing. Alice Lam: Investigation, Writing—review and editing. Scott R Manalis: Resources, Supervision, Funding acquisition, Project administration, Writing—review and editing.

## SUPPLEMENTARY FIGURES

**Figure 1–figure supplement 1.**
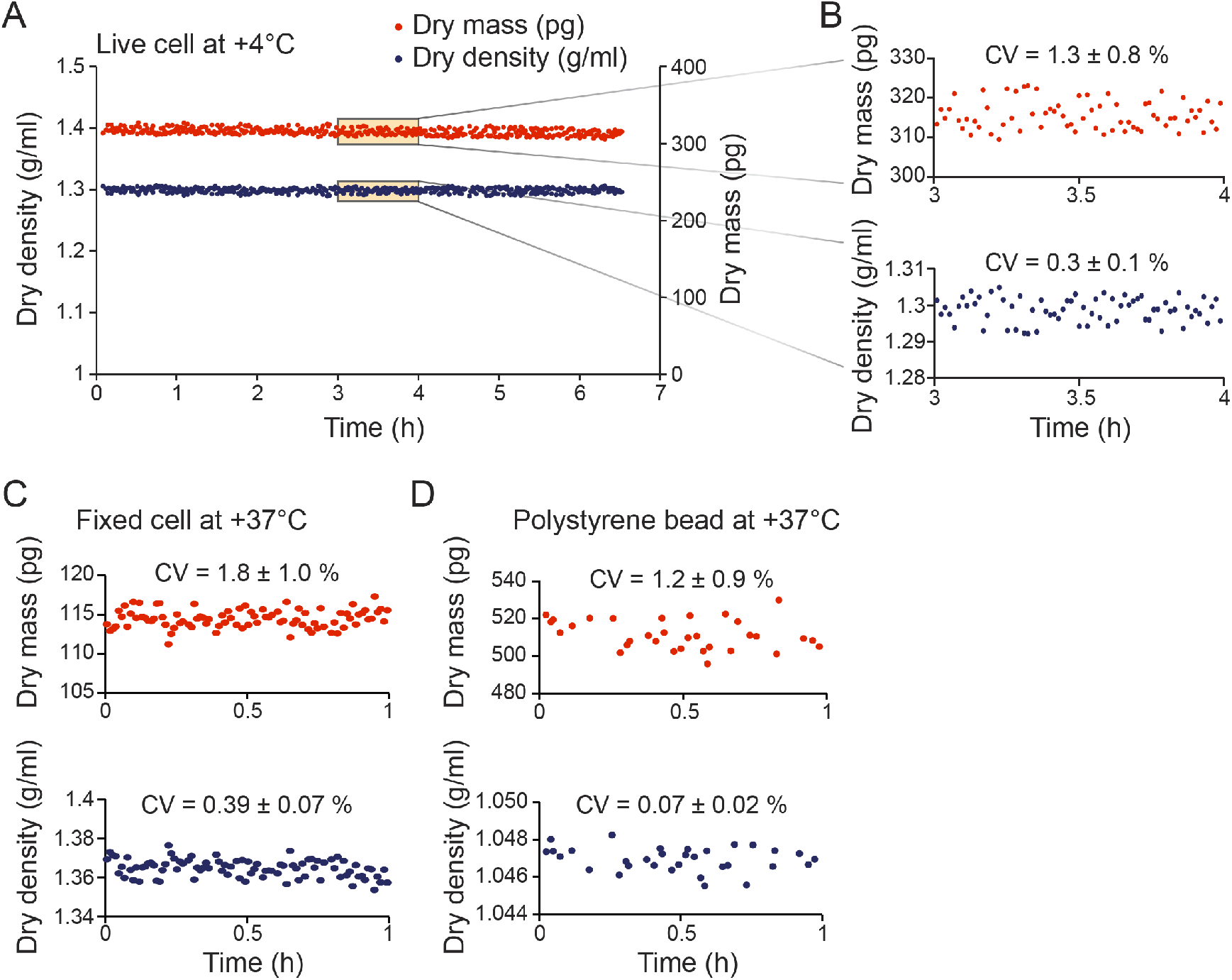
Quantifications of dry mass and dry density measurement errors. **(A)** Monitoring dry mass (red) and dry density (blue) of a single L1210 cell kept at +4°C to prevent cell growth and major compositional changes. **(B)** Zoom-ins of dry mass and dry density within the indicated one-hour window in panel (A). Measurement error was quantified as coefficient of variation (CV) within the one-hour window (mean±SD, N=6 independent experiments). **(C)** Monitoring dry mass and dry density of a single L1210 cell that was fixed to prevent cell growth and compositional changes. Measurement error was quantified as CV (mean±SD, N=5 independent experiments). **(D)** Monitoring dry mass and dry density of a 10 μm diameter polystyrene bead. Measurement error was quantified as CV (mean±SD, N=4 independent experiments). For all measurement error quantifications, the error in dry volume measurements was comparable to the error in dry mass measurements.

**Figure 1–figure supplement 2.**
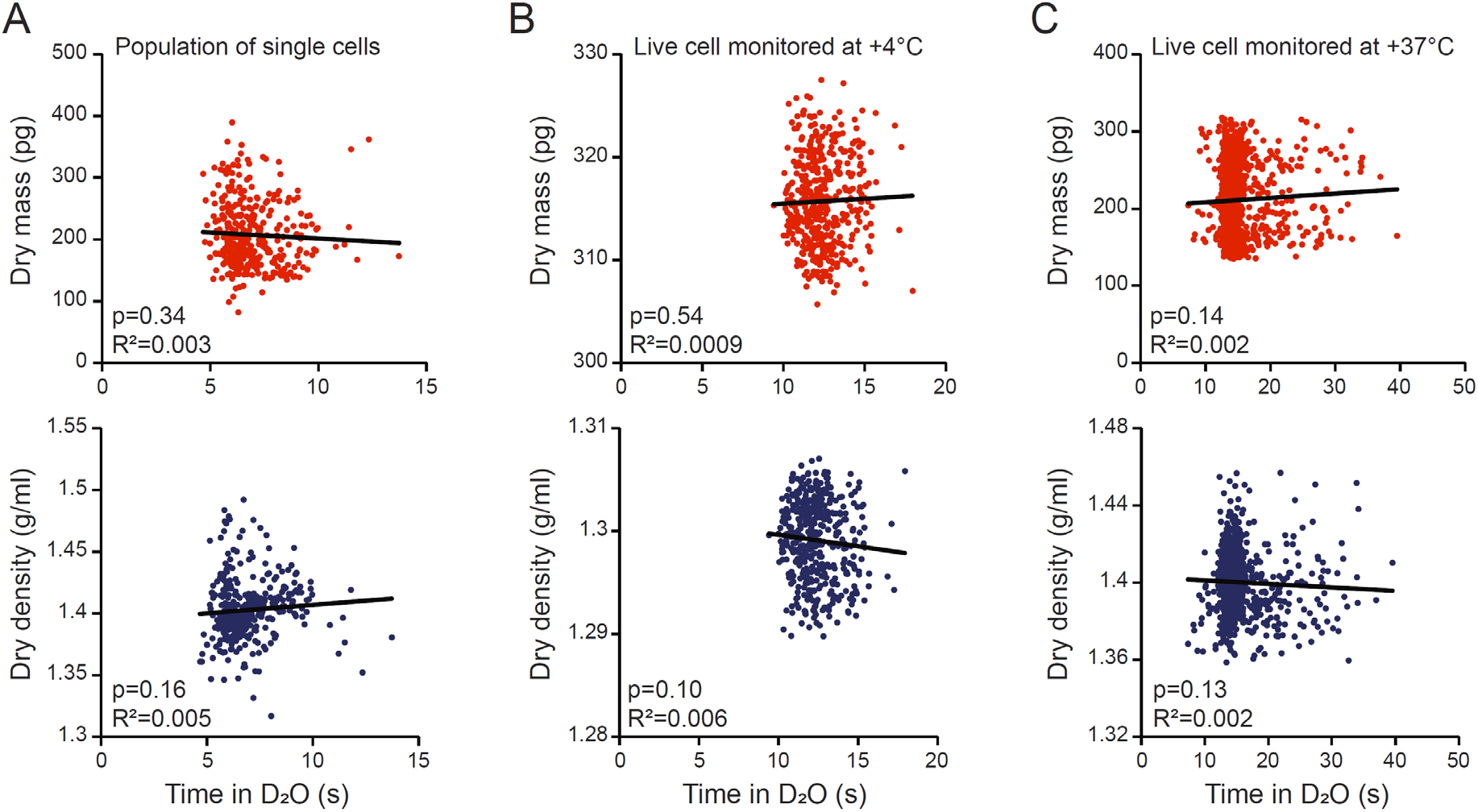
Dry mass and dry density measurements are not limited by the rate of water exchange. (**A**) Dry mass (top, red) and dry density (bottom, blue) as a function of time that the cell was exposed to D_2_O-based media between the two consecutive buoyant mass measurements. Each dot represents a single cell from a population of L1210 cells; n=358 cells. **(B)** Same as (A), but data is from single-cell monitoring of a L1210 cell at +4°C. Sama data as in Figure 1–figure supplement 1A; n=438 individual measurements. **(C)** Same as (A), but data is from single-cell monitoring of a L1210 growing at +37°C; n=1102 individual measurements. Each figure shows a linear fit, Pearson correlation (R^2^) and correlation p-values calculated using ANOVA.

**Figure 2–figure supplement 1.**
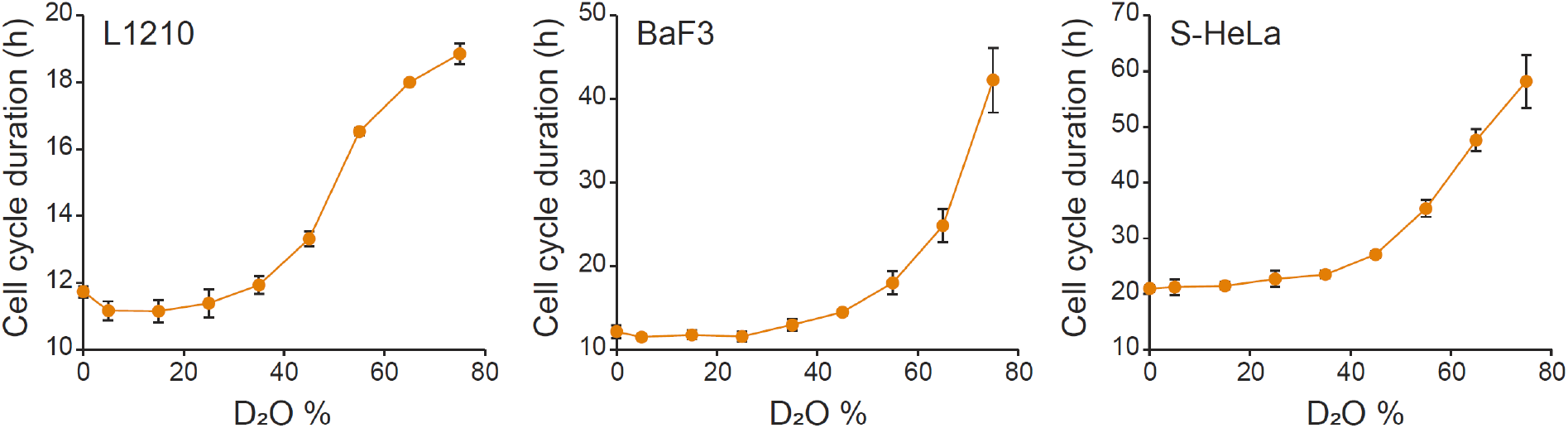
Influence of D_2_O on cell cycle durations in normal culture. L1210, BaF3 and S-Hela cells were grown in media, where indicated % of H_2_O was replaced with D_2_O. Cell proliferation was monitored by imaging cell confluency over ∼70h duration. N=5 independent cultures (mean±SD).

**Figure 2–figure supplement 2.**
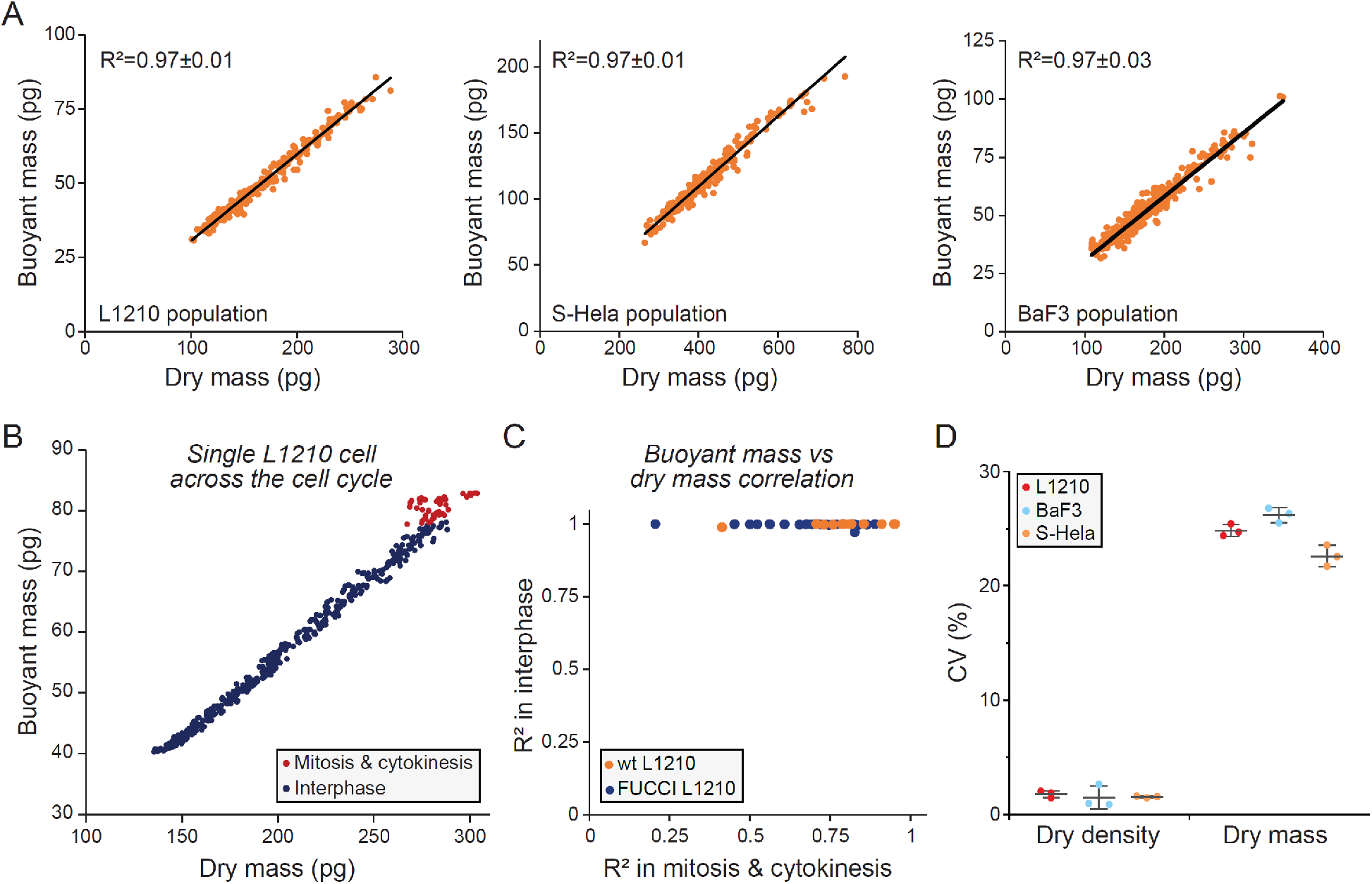
Dry and buoyant mass correlate near-perfectly, except in mitosis. (**A**) Correlation between dry and buoyant mass in L1210 (left), S-Hela (middle) and BaF3 (right) cells within a population. Each dot represents a single cell from the same experiment and black line is the linear fit. The Pearson correlation (R^2^) values represent the mean±SD from three independent experiments. (**B**) Correlation between buoyant mass and dry mass within a single wt L1210 cell cycle. Each dot represents a single measurement from the same cell in interphase (blue) or mitosis and cytokinesis (red). (**C**) Pearson correlations (R^2^) between buoyant mass and dry mass shown for interphase and for mitosis & cytokinesis. Data is obtained by single-cell monitoring and only full cell cycles are used (as in panel (B)). Each dot represents a single cell; data is shown for wt (orange) and FUCCI (blue) L1210 cells (n=12 and 27, respectively). (**D**) Cell-to-cell variability in dry density and dry mass in L1210 (left), S-Hela (middle) and BaF3 (right) cells within a population. Each dot represents an independent experiment (N=3); line and whiskers represent mean±SD.

**Figure 3–figure supplement 1.**
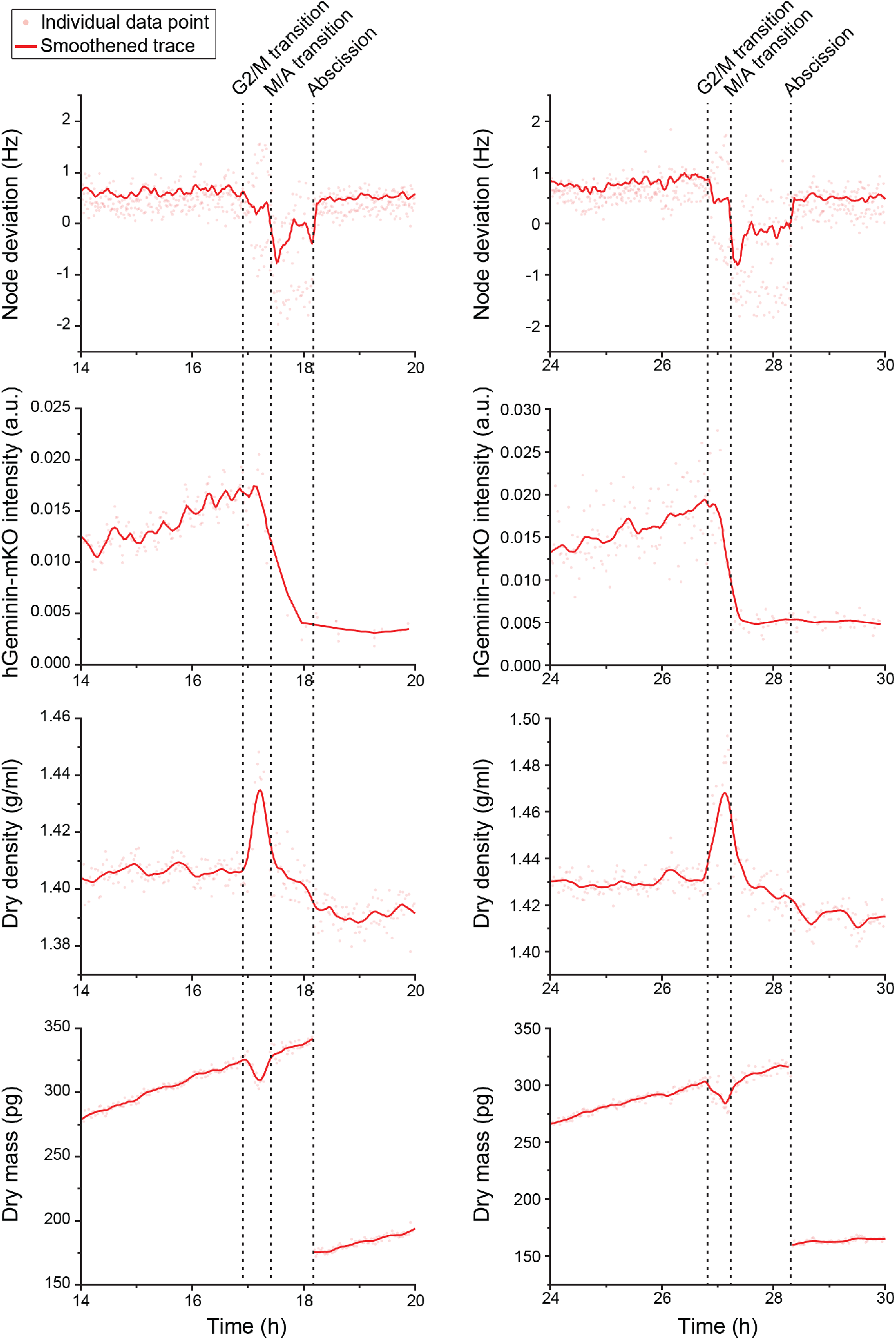
Cell cycle indicators & mitotic dry composition change. Two representative L1210 FUCCI cell traces (left and right column) with cell cycle indicators (top two rows) and dry composition changes (bottom two rows) in mitosis. Acoustic scattering from the cell, i.e. node deviation, is used to detect approximate mitotic entry and metaphase-anaphase transition (M/A transition), and the FUCCI cell cycle sensor, hGeminin-mKO fluorescence intensity, is used to validate the M/A transition.

**Figure 3–figure supplement 2.**
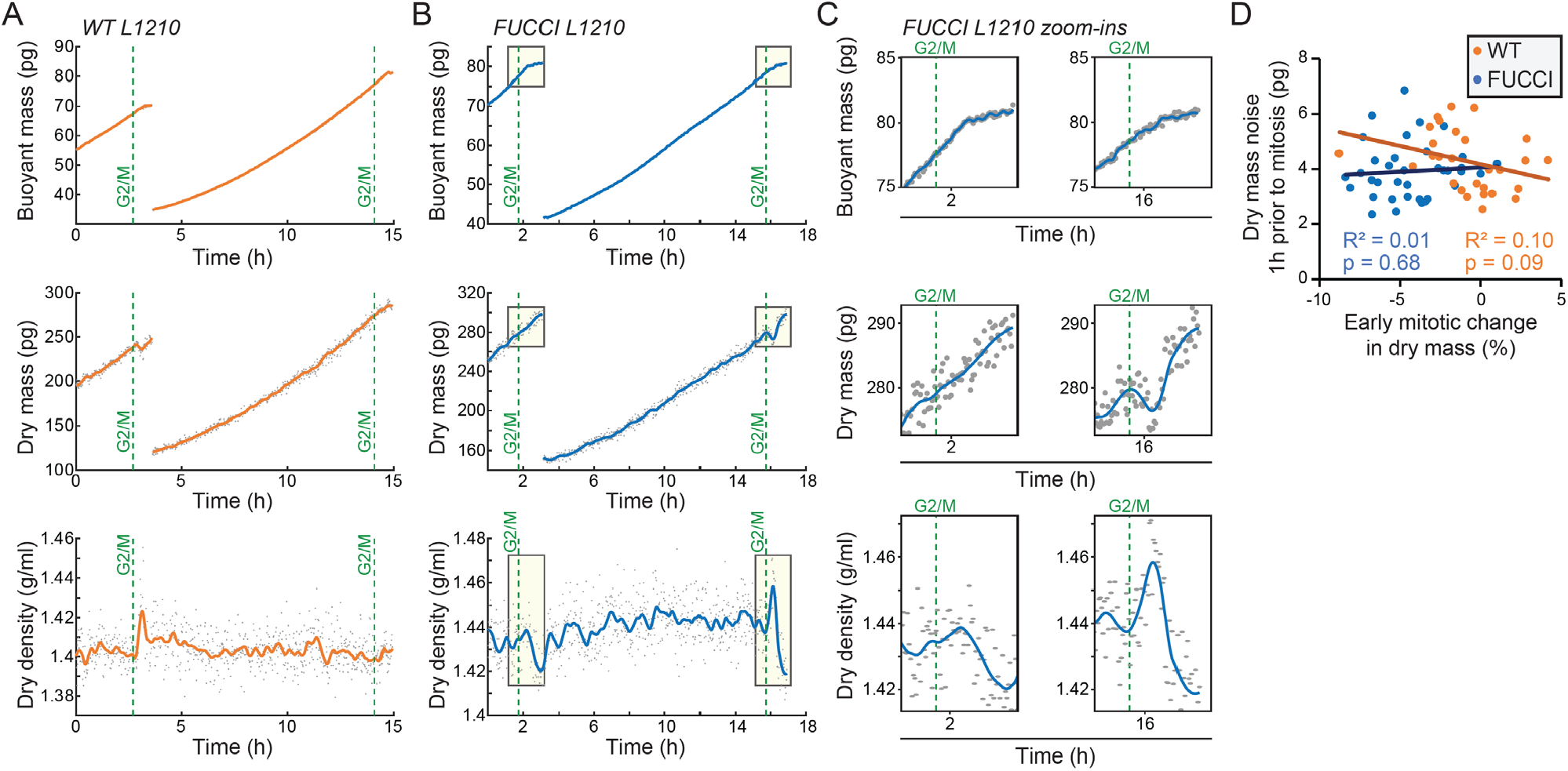
Cell-to-cell variability in the loss of dry mass in early mitosis reflects biological, non-genetic cell-to-cell variability. **(A)** Example buoyant mass (in H_2_O-based media), dry mass and dry density traces of wt L1210 cells, where mother and daughter cells display different mitotic dry composition behaviors. Grey dots indicate individual measurements and orange line indicates the smoothened trace; green, dashed vertical lines indicate mitotic entry (G2/M). **(B)** Same as panel (A), but for FUCCI L1210 cells. **(C)** Zoom-ins of the mitotic sections indicated by light yellow boxes in panel (B), showing the loss of dry mass and increase in dry density in daughter cell, but not the mother cell. Note that the buoyant mass resolution is lower than in our previous work (Miettinen et al., 2019) due to H_2_O and D_2_O fluid mixing which causes fluctuations in SMR baseline signal when monitoring the dry composition of a cell. **(D)** Correlation of experiment specific dry mass noise (deviation from a linear fit in the last 1h prior to mitotic entry) as a function of the early mitotic change in dry mass. Each point is a separate cell; correlation p-value calculated using ANOVA; N=28 cells for wt L1210; N=36 cells for FUCCI L1210.

**Figure 3–figure supplement 3.**
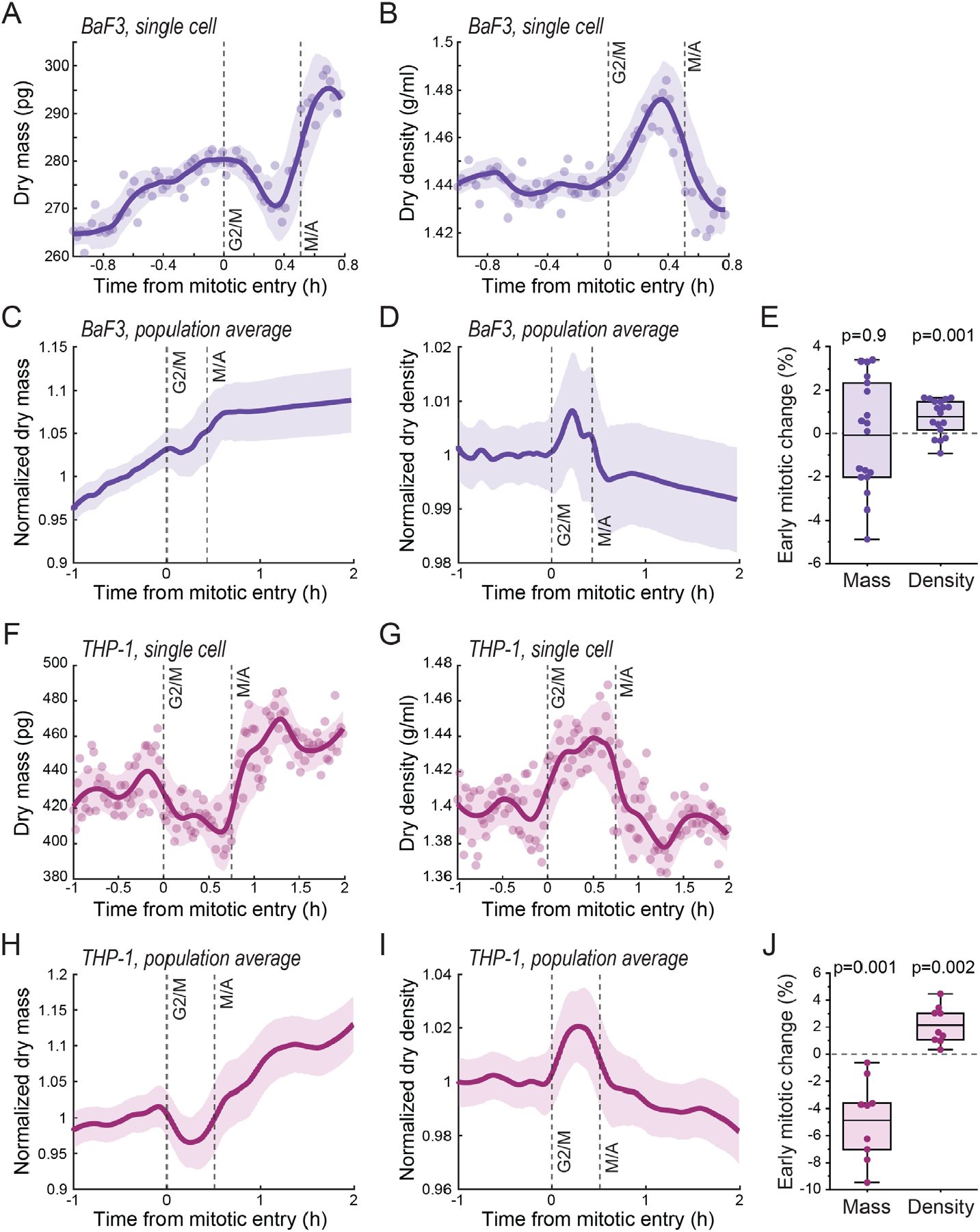
Mitotic dry mass and dry density behavior in BaF3 and THP-1 cells. **(A)** An example of BaF3 cell dry mass behavior in mitosis. Opaque points represent individual measurements; Thick line and shaded area represent smoothened data (mean±SD). Dashed vertical lines indicate approximate G2/M and metaphase-anaphase (M/A) transitions. **(B)** Same as (A), but for dry density. **(C)** Population average dry mass behavior in BaF3 cells. Thick line and shaded area represent mean±SD. N=18 cells from 11 independent experiments. **(D)** Same as (C), but for dry density. **(E)** Quantifications of dry mass and dry density changes in early mitosis in BaF3 cells. Dots represent individual cells; horizontal line represents mean; data is same as in panels (C&D); p-values calculated using one sample t-test and represent difference from zero. **(F-J)** Same as panels (A-E), but data is for THP-1 cells. N=9 cells from 8 independent experiments.

**Figure 4–figure supplement 1.**
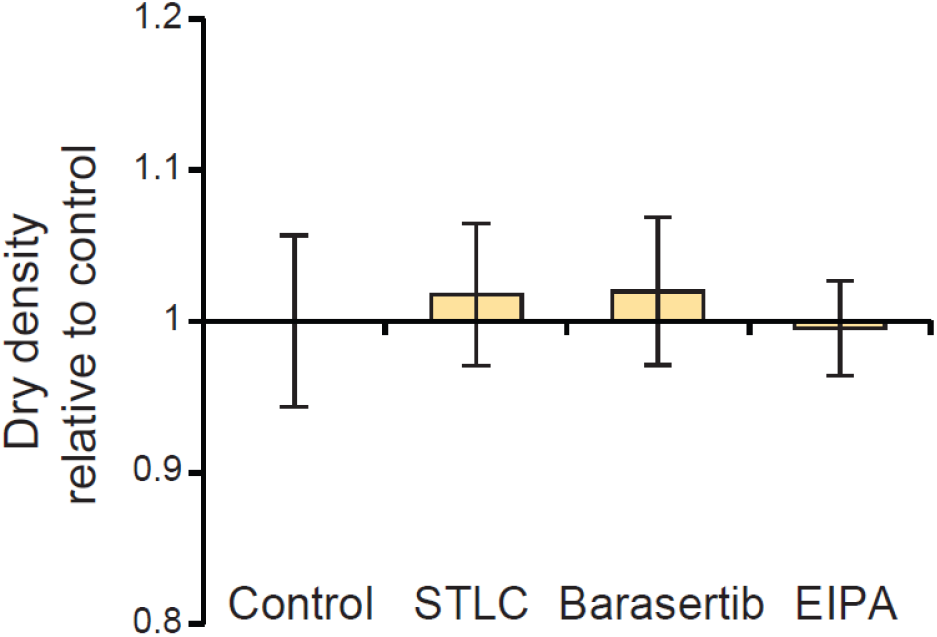
L1210 cell population dry density following mitotic perturbations. L1210 cells were treated with indicated chemicals for 4 hours using same concentrations as in Figure 4. Data represents mean±SD of dry density from >60 cells per condition.

**Figure 5–figure supplement 1.**
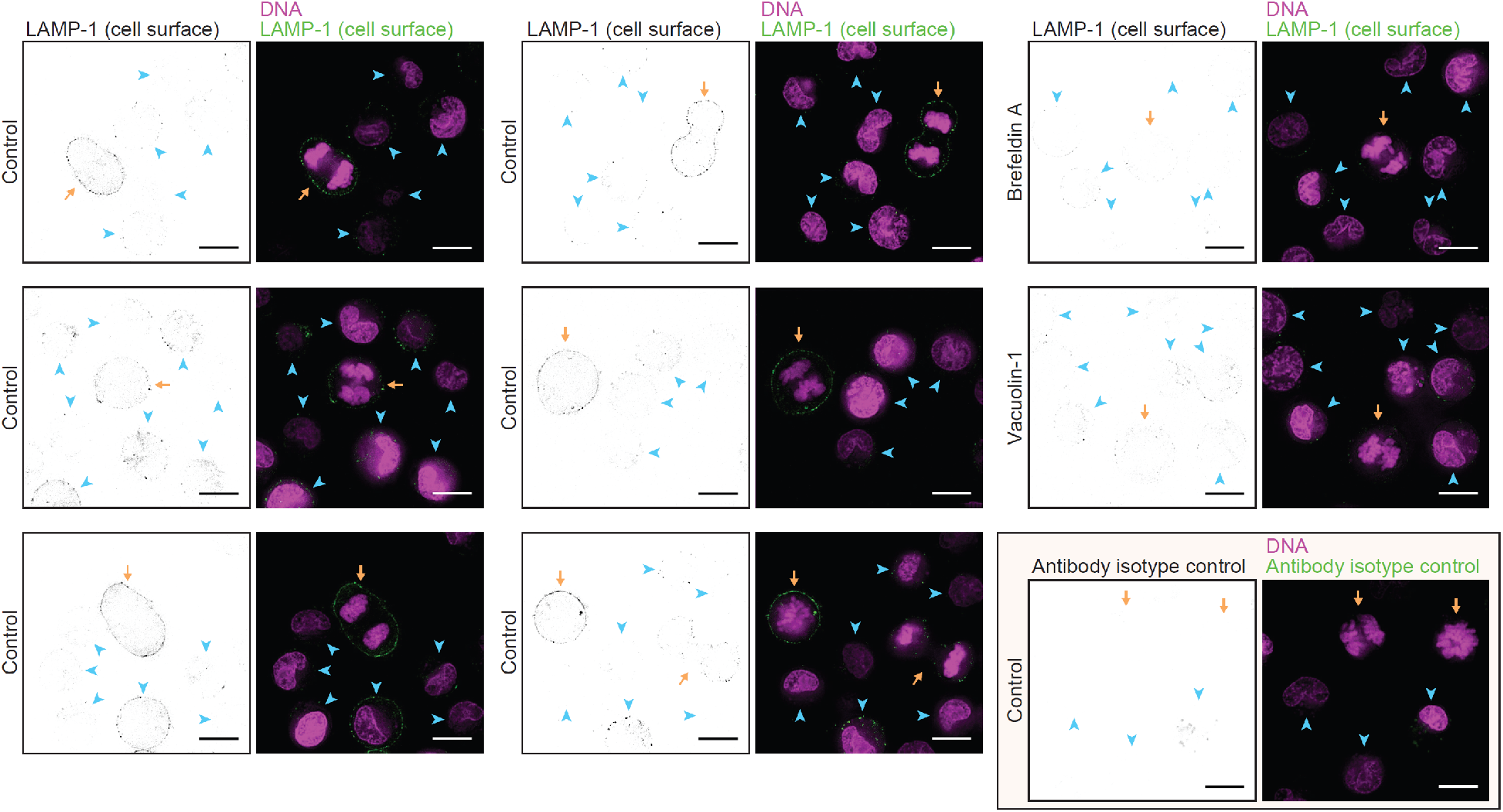
Additional examples of live L1210 cell surface LAMP-1 labeling. Additional images of surface LAMP-1 immunolabeling alone (left, black&white) and together with DNA labeling (right, color) in live L1210 cells. First two columns display LAMP-1 immunolabeling under control conditions. Last column displays LAMP-1 immunolabeling after 4h drug treatments (top two rows) or under control conditions but using the corresponding antibody isotype labeling (bottom row). All images are from wt L1210 cells and represent a single z-slice. Orange arrows indicate mitotic cells, blue arrowheads indicate interphase cells. All scalebars depict 10 μm.

**Figure 5–figure supplement 2.**
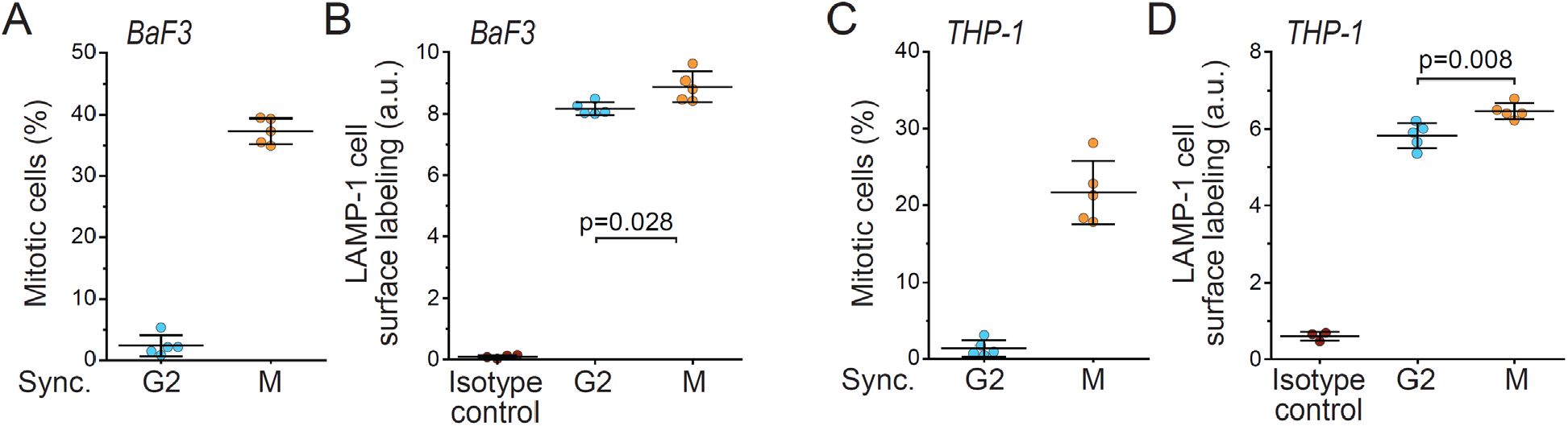
Lysosomal exocytosis is higher in early mitosis than in G2 in BaF3 and THP-1 cells. **(A)** % of mitotic cells in BaF3 cell populations following synchronization to G2 or early mitosis. N=5 independent cultures. **(B)** Flow cytometry quantifications of BaF3 cell population surface LAMP-1 immunolabeling following synchronization to G2 or early mitosis. N=4-5 independent cultures. Isotype control indicates labeling intensity when using isotype control antibody in an unsynchronized population. **(C)** Same as panel (A), but for THP-1 cells. N=5 independent cultures. **(D)** Same as panel (B), but for THP-1 cells. N=3-5 independent cultures. In all figures, line and whiskers indicate mean±SD, p-values calculated using Welch’s t-test.

**Figure 6–figure supplement 1.**
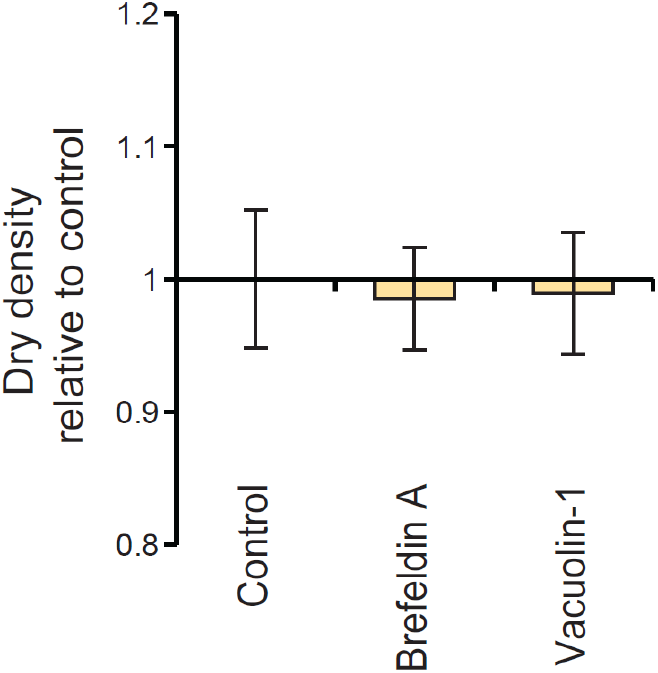
L1210 cell population dry density following exocytosis perturbations. L1210 cells were treated with indicated chemicals for 4 hours using same concentrations as in Figure 6. Data represents mean±SD of dry density from >70 cells per condition.

